# Modeling integration site data for safety assessment with MELISSA

**DOI:** 10.1101/2024.09.16.613352

**Authors:** Tsai-Yu Lin, Giacomo Ceoldo, Kimberley House, Matthew Welty, Thao Thi Dang, Denise Klatt, Christian Brendel, Michael P. Murphy, Kenneth Cornetta, Danilo Pellin

**Affiliations:** National Gene Vector Biorepository, Department of Medical and Molecular Genetics, Indiana University School of Medicine, Indianapolis IN, USA; Division of Hematology/Oncology, Boston Children’s Hospital, Harvard Medical School, Boston, MA, USA; Department of Pediatric Oncology, Dana-Farber Cancer Institute, Harvard Medical School, Boston, MA, USA; Department of Surgery, Indiana University School of Medicine, Indianapolis IN, USA; Harvard Data Science Initiative, Cambridge, MA, USA

## Abstract

Gene and cell therapies pose safety concerns due to potential insertional mutagenesis by viral vectors. We introduce MELISSA, a statistical framework for analyzing Integration Site (IS) data to assess insertional mutagenesis risk. MELISSA employs regression models to estimate and compare gene-specific integration rates and their impact on clone fitness. We tested MELISSA under three settings. First, we conducted simulation studies to verify its performance under controlled conditions. Second, we characterized the IS profile of a lentiviral vector on Mesenchymal Stem Cells (MSCs) and compared it with that of Hematopoietic Stem and Progenitor Cells (HSPCs), in addition to comparing the in vitro clonal dynamics of MSCs isolated from alternative tissues. Finally, we applied MELISSA to published IS data from patients enrolled in gene therapy clinical trials, successfully identifying both known and novel genes that drive changes in clone growth through vector integration. MELISSA identifies over– and under-targeted genes, estimates IS impact and explores biological relevance through pathway analysis. This offers a quantitative tool for researchers, clinicians, and regulators to bridge the gap between IS data and safety and efficacy evaluation, facilitating the generation of comprehensive data packages supporting Investigational New Drug (IND) and Biologics License (BLA) applications and the development of safe and effective gene and cell therapies.

## Introduction

Gene therapy using hematopoietic stem and progenitor cells (HSPC) modified with viral vectors has emerged as a promising approach for treating rare monogenetic diseases (Ferrari et al., 2021; Naldini, 2015). This strategy involves introducing functional copies of therapeutic genes into patients’ HSPCs, enabling them to produce the missing or malfunctioning protein and restore normal cellular function. Due to their efficient gene delivery capabilities, viral vectors have also found extensive application in novel cell therapies such as Chimeric Antigen Receptor (CAR)-T immunotherapies, where Lenti/Retroviral Vectors are used to deliver CAR genes in a patient’s T cell.

The capability for viral vectors to integrate their cargo DNA into unpredictable locations within the host genome carries a certain risk of Insertional Mutagenesis (IM), defined as the disruption or dysregulation of a gene caused by a vector insertion. IM can lead to enhanced cell growth due to the activation of oncogenes or the disruption of tumor suppressor genes by the integrated DNA (Baum et al., 2004; Bushman, 2020; Cornetta et al., 2023; Woods et al., 2006). Clonal dominance underscores the risk of a particular clone or a small subset of clones of transduced cells gaining a growth advantage, potentially skewing the clonal composition of the repopulating cells towards monoclonal or oligoclonal configurations that pose additional long-term safety and efficacy concerns, including oncogenic transformation (Corre & Galy, 2023; Fehse & Roeder, 2008; Nienhuis et al., 2006).

Regulatory agencies like the US FDA require rigorous preclinical safety and efficacy evaluations and 15 years of IM monitoring for patients treated with genetically modified HSPCs (Fda & Cber, 2020). Recently, the US FDA extended monitoring to individuals receiving BCMA-or CD19-directed autologous CAR T cell immunotherapies. A comprehensive risk-benefit analysis is crucial for evaluating the potential for IM, yet the scientific community still lacks standardized indices, metrics, and methods to clearly distinguish between safe and unsafe integration profiles.

The association of the integrome, namely the distribution of IS across the host genome, with genomic annotation data has elucidated the role of chromatin conformation, transcriptional activity, and the cellular genome 3D nuclear organization on IS frequency (Biasco et al., 2011; Cattoglio et al., 2010; Goff, 2007; Hofmann et al., 2006; Lelek et al., 2015). These insights, combined with the analysis of oncogenic transformation mechanisms observed in clinical trials, enabled the generation of novel and safer viral vectors for HSPC gene therapies.

Using IS analysis to monitor the clonal composition over time, scientists can gain insights into the diversity and dynamics of the engrafting cell population, assess the long-term efficacy, and the risk of clonal dominance (Corre & Galy, 2023; Pellin et al., 2023; Scala et al., 2018; Six et al., 2020). From a safety perspective, IS analyses serve as the primary screening tool for identifying potential risks associated with uncontrolled clonal expansion by detecting abundant clones based on their relative contribution and mapping IS location within the host cell genome. Conventionally, the identification of abundant clones has been based on predefined percentage thresholds calculated on an individual sample basis.

There is currently no method to estimate integrome and clonal contribution using information from multiple time points, cell types, or patients, which would allow for more accurate characterization of gene targeting preferences and detection of clonal expansion. This paper introduces a novel set of statistical tools, MELISSA (ModELing IS for Safety Analysis), to translate IS data into actionable safety assessment and evaluation insights. MELISSA provides statistical models for measuring and comparing gene targeting rates and their effects on clone growth within a gene-based approach. MELISSA modeling consists of a regression approach that analyzes and combines data from complex experimental designs, including datasets with multiple patients or donors, replicates, and additional covariates of interest. MELISSA facilitates the quantitative comparisons of different conditions and includes rigorous statistical testing, visualization, and annotation to aid the biological interpretation of results.

From an experimental standpoint, we constructed a unique dataset to compare the integration and safety profiles of Mesenchymal Stem Cells (MSCs) isolated from two different sources and mobilized CD34+ Hematopoietic Stem and Progenitor Cells (HSPCs). This enabled us to evaluate the effectiveness of our proposed framework as a tool for analyzing preclinical experiments and guiding the development of novel gene therapy strategies.

Additionally, we analyzed published IS data from patients enrolled in gene therapy clinical trials for Beta-thalassemia (β-thal), Sickle Cell Disease (SCD), and Wiskott-Aldrich Syndrome (WAS) (Six et al., 2020). Our analysis revealed that, although none of the clones in the available IS data reached a relative abundance considered concerning (max single clone relative abundance: β-thal, 1%; SCD, 2.4%; WAS 9.5%), by leveraging longitudinal modeling of clone sizes, we identified both known and novel genes with the potential to affect clone fitness when targeted by an IS. This highlights MELISSA’s sensitivity and ability to detect early signs of clonal expansion, demonstrating its effectiveness in identifying risk factors before they become prominent.

MELISSA is a quantitative tool for supporting researchers, clinicians, and regulatory agencies in advancing the development of safe and effective gene and cell therapies. Its goal is to bridge the gap between the IS datasets generated from preclinical experiments to support IND applications and clinical samples for safety and efficacy evaluation.

## Results

MELISSA is a robust statistical framework available as an R package designed to analyze and interpret IS data for safety evaluation in viral vector-based gene and cell therapies. We evaluated MELISSA using a comprehensive Monte Carlo simulation study to assess performance metrics, specifically focusing on the Positive Predictive Value (PPV) and the detection rate. This allowed us to understand the proportion of true positive results among all positive test results, reflecting the reliability of a positive test outcome and the likelihood of detecting a real effect or difference when it exists.

As shown in the schematic in Figure 1, it requires three primary inputs. The IS tables are provided in bed file format and contain clone size estimates (as read counts, UMIs, or shear site data) (Aird et al., 2011; Berry et al., 2012; Brady et al., 2011). The design matrix includes sample-specific covariates, such as conditions, replicates, cell type, time after therapy, and others. MELISSA can analyze the full set of genes using genome annotation files, providing a genome-wide comprehensive overview of potential IM risks. Alternatively, it allows users to define specific regions of interest within the genome for targeted analysis.

**Figure 1:**
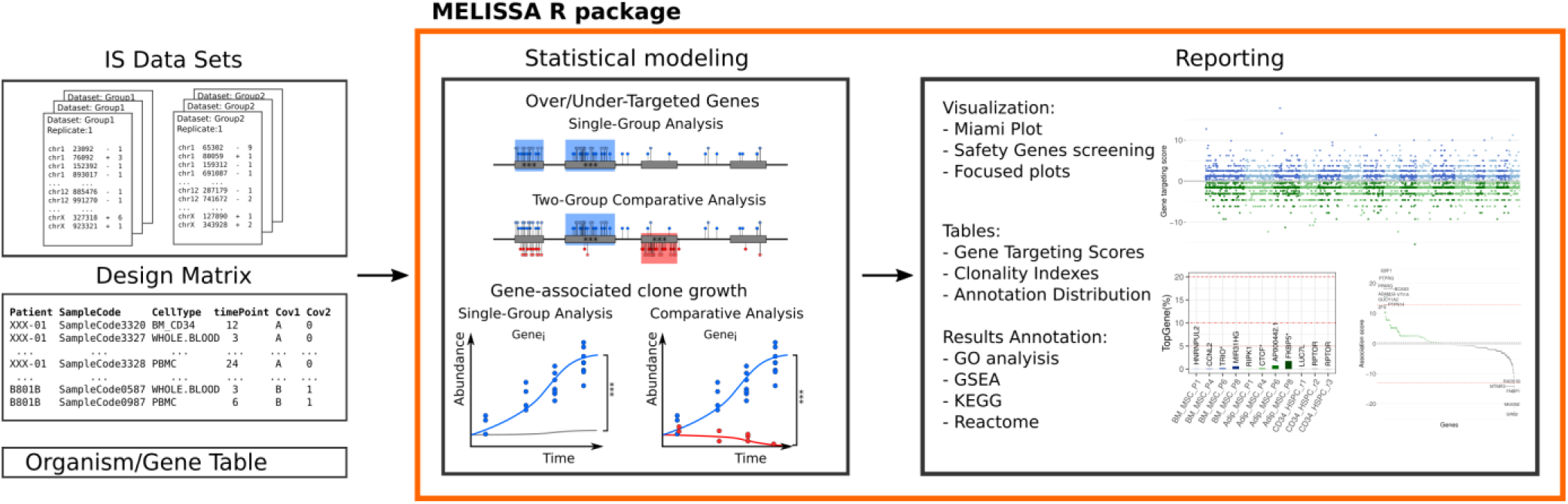
Schematic of MELISSA workflow. MELISSA receives as input 1. IS data in a bed file format, 2. a design matrix with all the relevant samples’ annotation, 3. a reference genome or a bed file containing genomic region to be tested for IS enrichment. MELISSA combines input data and performs two main types of analysis: gene targeting rates and gene/clone fitness alterations. MELISSA also has a dedicated function for calculating clonality indexes, graphical representation of the results, and downstream pathway enrichment analyses.

MELISSA provides a comprehensive suite of functionalities. It generates descriptive statistics summarizing IS distribution across various genomic features and calculates samples’ clonality indexes commonly referenced in the literature. Additionally, it tackles critical safety-related questions: 1. estimating gene-specific IS rates within a single group or comparing them across groups to identify potential differences across conditions; 2. estimating whether IS within a gene influences clone growth, again for single or multiple groups.

Furthermore, MELISSA facilitates downstream analysis through various methods. Researchers can generate informative graphical representations and utilize gene-scoring tables to identify potential associations between IS data and underlying biological mechanisms with tools such as Gene Ontology (GO) enrichment analysis, Gene Set Enrichment Analysis (GSEA) and KEGG and REACTOME pathway analysis (Ashburner et al., 2000; Fabregat et al., 2017; Kanehisa et al., 2017). Additionally, the distribution of genomic annotations, provided in BED file format (e.g., epigenetic profiles) within IS flanking regions, offers deeper insights into the mechanisms driving IS selection process.

### Statistical modeling

For estimating and comparing gene-specific IS rates, we developed a logistic regression model to identify over– and under-targeted genes and calculate gene-specific targeting rates (Methods). This approach considers the presence/absence of IS at each genomic coordinate as a binary variable (1 if an IS maps to the specific location and 0 otherwise). In the single-condition setting, the likelihood of an IS event occurring in a specific gene is compared to the remaining part of the genome. The gene IS-enrichment score is calculated through a regression parameter that quantifies the ratio between the gene’s IS rate and the baseline, genome-wide IS rate. This model intrinsically normalizes datasets for factors such as gene length and dataset size. The two-condition differential analysis is specifically designed to allow IS rates to vary across genes, and our detecting strategy will highlight only genes with an unshared variation of the gene-targeting rates across conditions.

The rationale for estimating the impact of IS on clone growth is based on the biological assumption that IS within (or in close proximity to) a specific gene might be able to perturb its expression profile, either through the action of the transgene’s promoter sequence, by the alteration of endogenous critical regulatory elements or other mechanisms (Bushman, 2020; Ranzani et al., 2013). By grouping and modeling the growth dynamics of all clones harboring IS within a specific gene, we aim to discriminate clone growth due to stochastic clonal selection from IS-driven mechanistic effects. Under this scenario, we model longitudinal clone size data using logistic regression for binomial (count) data and detect groups of gene-associated IS that significantly expand faster over time using a statistical test (Methods). Comparing differential gene/clone growth analysis allows us to estimate whether IS in the gene under investigation affects clone growth differently in the two conditions.

In all the scenarios described, we use Likelihood Ratio Tests (LRT) to assess gene associations, with p-values adjusted using the False Discovery Rate (FDR) (Benjamini & Hochberg, 1995) or Holm– Bonferroni (Holm, 1979) methods. The LRT statistic serves as a score for gene targeting or gene-clone growth associations and, in differential analysis, is multiplied by the sign of the estimated parameter to reflect the direction of the effect.

### Simulation study

To evaluate MELISSA’s performance in detecting variations in IS targeting rates and gene-specific effects on clone fitness, we conducted a simulation study on chromosome 15 (hg38) in which we included five artificial genes (*TestGenes*) with a length of 5, 10, 20, 40, and 80 kb pairs not overlapping with the existing 1027 endogenous genes. The study assessed how the PPV and *TestGenes’* detection rate varied with sample size and effect strength (Figure 2, Suppl. Table 1). For each setting, 1,000 simulations are used to assess the model’s performance. Further details of the simulation design are provided in the Methods section, and the simulated datasets are shown in Suppl. Figure 1.

**Figure 2:**
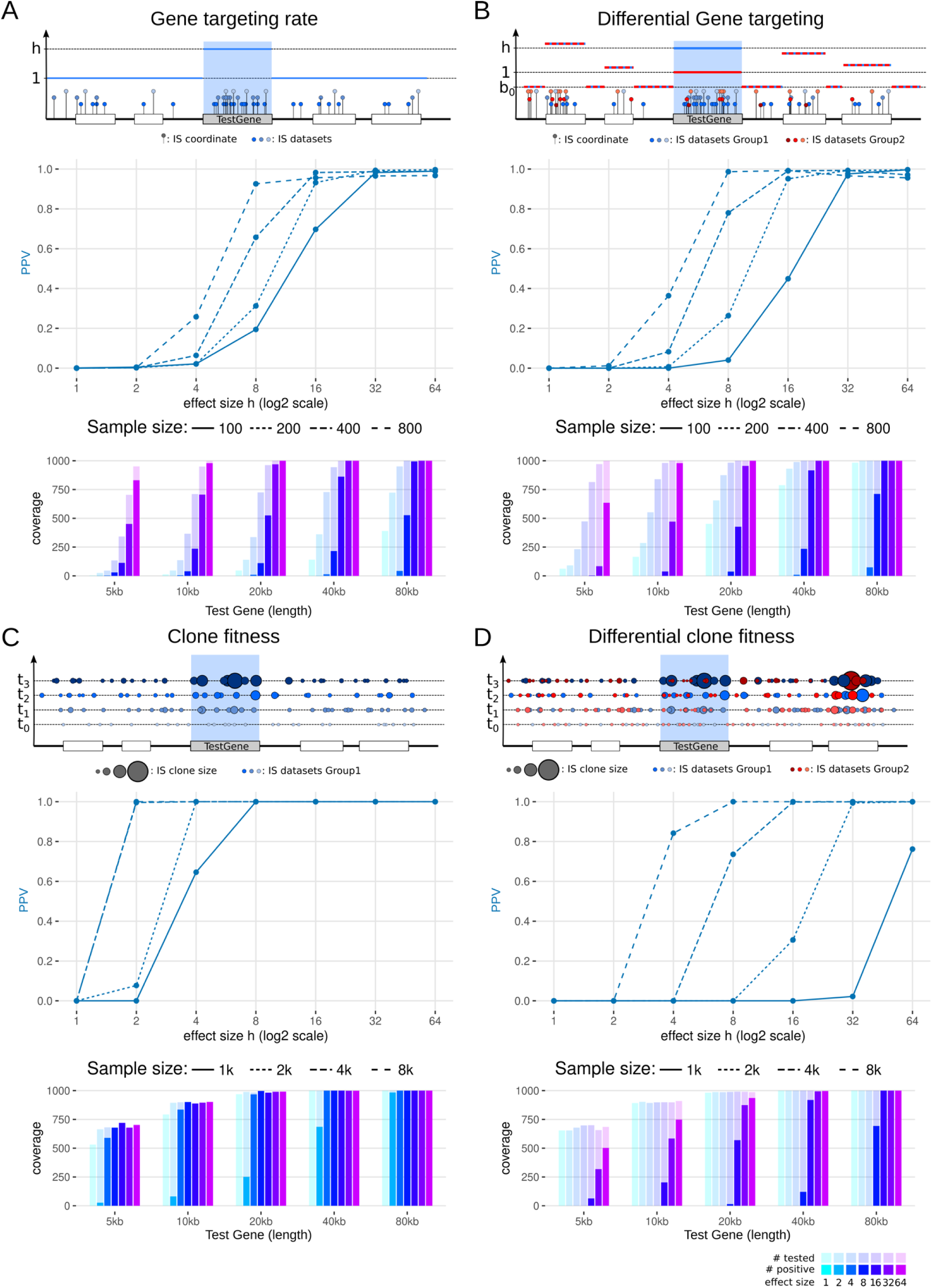
Simulation study results. Average Positive Predicted Value (PPV) across 1000 simulations for model A) gene targeting rate; B) differential gene targeting; C) clone fitness; D) differential clone fitness. Solid, dashed, dotted, and dash-dot lines correspond to different sample sizes. Seven increasing effect sizes have been tested, ranging from h=1 (no effect) to 64. Detection rate bar graphs (bottom of each panel) report the number of times each *TestGene* has been tested (semi-transparent bars) and the number of times in which the *TestGene* is detected as significantly over-targeted/gene growth associated, for the simulations with sample size N=400 for A), B), and M=4000 for C), D)

Two strategies were used to simulate IS datasets for gene targeting rate analysis. In the single-group analysis (Figure 2A), *N* = 100, 200, 400, 800 IS elements were generated, corresponding to genome-wide IS datasets ranging from 3,000 to 25,000 IS, in line with preclinical and clinical datasets. IS were simulated using a two-part model: a genome-wide baseline and elevated integration rates for over-targeted genes, with fold increases of *h* = 1 (no effect), 2, 4, 8, 16, 32, and 64. For differential targeting analysis (Figure 2B), each gene was assigned a unique rate, consistent across datasets, with group 1 showing elevated integration probabilities compared to group 2.

To analyze clone fitness and detect variations in clone growth rates, we simulated time-series data of IS clone sizes across six time points (*t* = 0 to 5). We first generated a database of 10,000 potential IS coordinates using a constant baseline and a variable gene-specific targeting rate. Clone sizes were determined by sampling IS from this database, with replacement, at each time point. Sampling probabilities for IS in *TestGenes* linearly increase over time (Figure 2C), mimicking selective clone expansion. In the single-group scenario, datasets with cumulative clone sizes of *M* = 1000, 2000, 4000, 8000 were generated. For differential analysis (Figure 2D), 12 datasets were generated, representing two groups with identical sampling probabilities, except for *TestGenes* where group 1 had higher growth rates controlled by the effect size *h*.

We evaluated the impact of p-value adjustment methods on the performance of MELISSA, with results shown in Suppl. Figure 2. Using adjusted p-values, MELISSA’s PPV consistently outperformed results obtained using raw p-values, emphasizing the importance of correction methods in improving predictive accuracy. The Holm and FDR adjustment methods yielded comparable results. Computational times under different scenarios are detailed in Suppl. Table 2.

Our simulation study demonstrated that MELISSA’s PPV and detection rates consistently improved with increasing sample sizes and effect strengths. In single-group analyses, detecting higher gene targeting rates in short genes proved challenging, while clone fitness analysis showed strong sensitivity, even for low effect sizes. For the differential setting, the results highlighted that detecting differences in gene targeting rates is more efficient than detecting variations in clone fitness. Importantly, the model produced no false positives, confirming its high specificity.

### Using MELISSA to evaluate clonal dynamics of transduced MSC over time

MSC is a heterogeneous cell population found in bone marrow, adipose tissue, placenta, umbilical cord and cord blood, dental pulp, and other tissues that show potential for cell therapy applications due to their regenerative potential. There are over 1500 clinical trials exploring their application across diverse diseases (www.clinicaltrial.gov), and lentiviral vector (LV) are being evaluated to increase the therapeutic potential of MSCs. While extensive research has been conducted on the integrome in other cell types, our understanding of LV integration and its potential risks in MSCs remains limited.

Low passage healthy donor Bone Marrow-derived (BM), Adipose-derived (Ad) MSCs, and CD34+ healthy donor mobilized HSPCs were transduced with an LV encoding EGFP. IS analysis was performed on all cell types early after transduction and at various passages of in-vitro culture for MSCs (P1, P4, P6, P8). Cells transduced at the initiation of the culture will give rise to a progeny of cells (IS clone) that share the exact IS coordinates (i.e. the LV IS). More details on the MSC transduction, IS retrieval protocol and bioinformatic analysis are available in the Methods section.

Accurate clone size estimation is crucial for IS analysis since it allows for understanding clones’ relative contribution and characterizing the composition of the population. Here we estimated clone sizes using the sonicLength method (Berry, 2014), albeit alternative quantification strategies (read counts, UMIs) are compatible with the MELISSA framework. As shown in Figure 3A and Suppl. Table 3, the cumulative clone size in BM MSCs shows a consistent positive trend throughout the in-vitro culture period. In contrast, the number of distinct clones steadily increases in P1 through P6, with a drop observed at P8. Ad MSCs displayed a declining IS count and a similar trend in total clone size. These observations are further supported by the Shannon diversity index analysis (Figure 3B), indicating stable diversity in BM MSCs and a progressive loss of diversity in Ad MSC.

**Figure 3:**
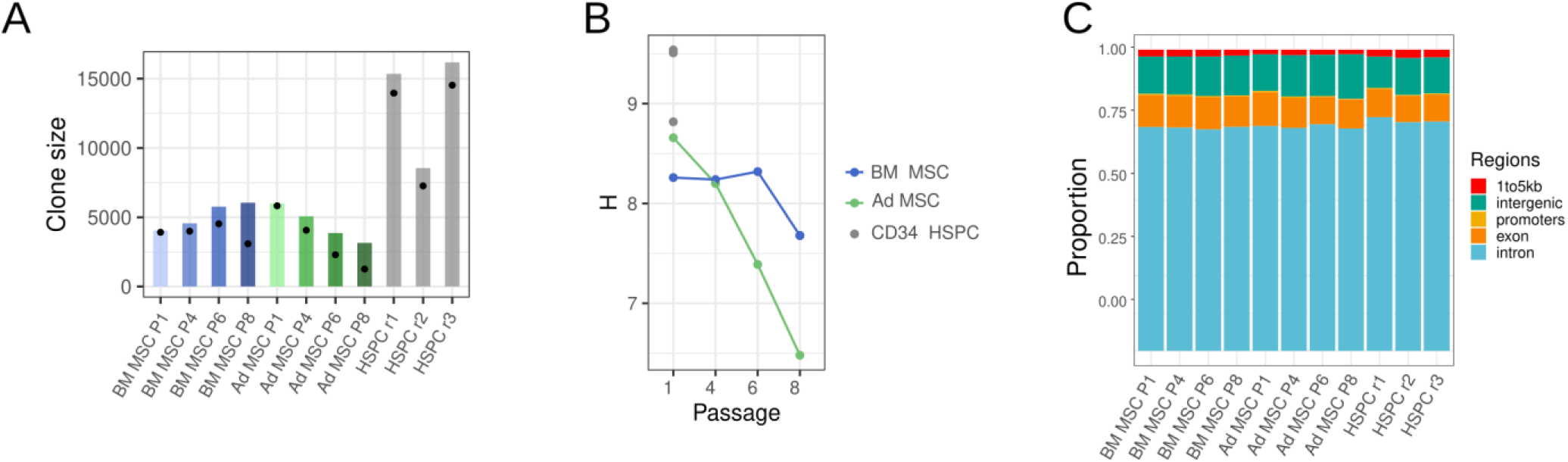
IS datasets descriptive statistics. A) Integration site counts and cumulative clone size: Bar chart depicting the cumulative clone size for each dataset. The number of distinct IS retrieved in each sample is reported with dots. B) Shannon entropy (H) index. Line plot representing the longitudinal dynamics of the diversity during in-vitro culture with subsequent passages of MSC. CD34 HSPC were analyzed at a single timepoint. C) Integration site annotation distribution: stacked bar chart displaying the distribution of integration sites according to genomic annotation, delineating the genomic regions where integration events are most prevalent.

LV integration preferences have been primarily reported in hematopoietic cells. The distribution observed in our HSPC datasets confirmed the known tendency of LV vectors to integrate into gene bodies. Interestingly, we found that this pattern also showed remarkable consistency in MSCs (Ambrosi et al., 2011; Cattoglio et al., 2007; Schröder et al., 2002). In the gene-based analysis reported in the following sections, we extended the gene bodies (region from the transcription start site to the end of the transcript) by including the 1-kb gene promoter region upstream of a gene’s transcriptional start site, thereby encompassing approximately 85.2% of the IS database (74.9% intronic, 10% exonic, and 0.3% resided in gene promoter regions).

### MELISSA identifies both shared and cell type-specific over-targeted genes in MSCs and HSPC

We analyzed IS datasets using MELISSA to identify the most targeted genes in BM MSC, Ad MSC and HSPC to investigate the presence of differentially targeted genes.

Figure 4A shows the gene scores from the HSPC control dataset analysis. Among 1959 significant genes, known Recurrent Integration Genes (RIGs) like *PACS1*, *KDM2A*, and *GRB2* have higher scores (Yan et al., 2023), confirming our approach effectively captures established integration patterns.

**Figure 4:**
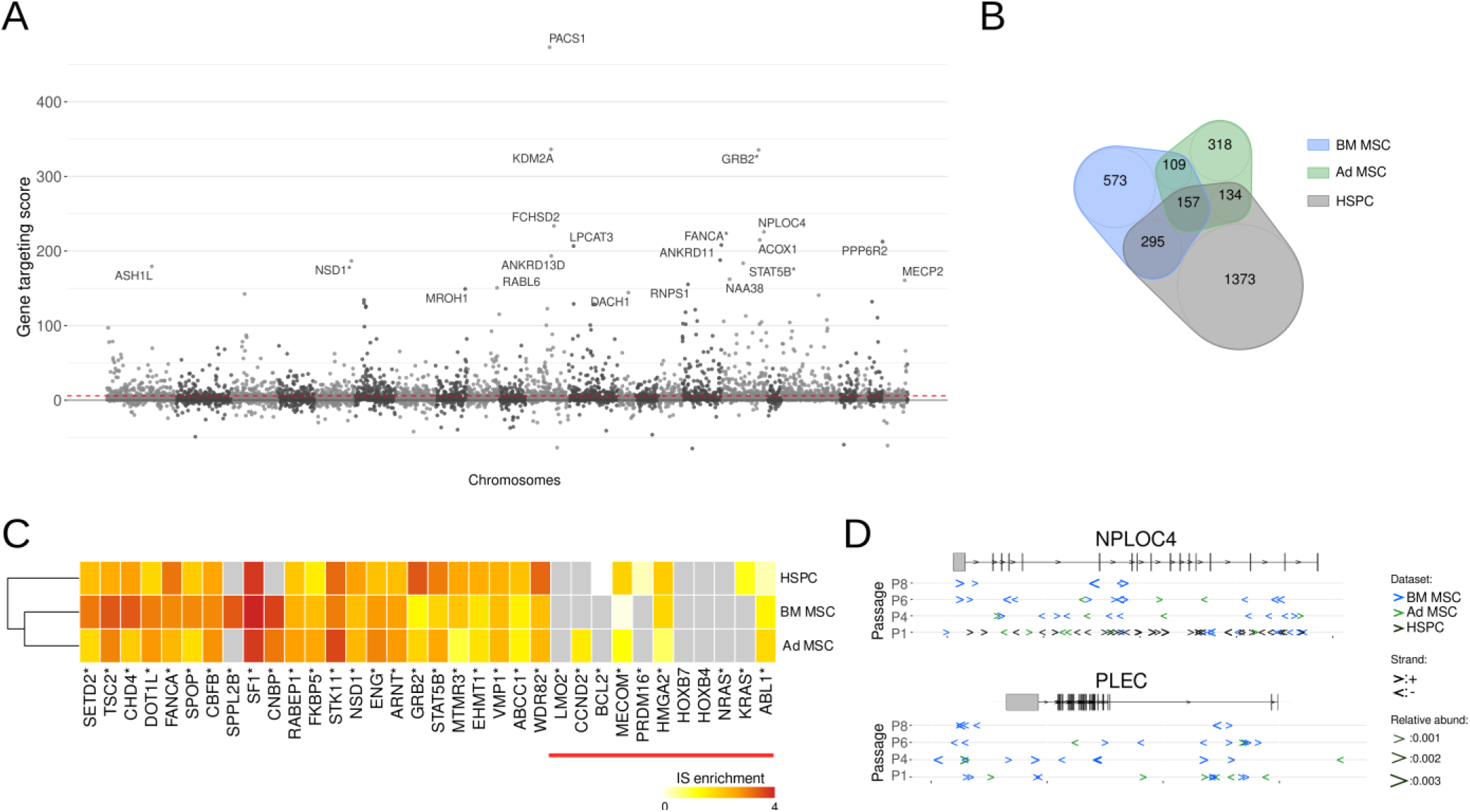
Results for the single group gene targeting rates analysis. A) Gene targeting score for the HSPC dataset. Each dot corresponds to a gene. On the x-axis, genes are ordered according to chromosome location, with genes in even-numbered chromosomes having a lighter color tone. The gene targeting score on the y-axis corresponds to signed LRT test statistics. Genes marked with a * are included in the high-risk gene list. The dashed red line represents the threshold for statistically significant enrichment (LRT p-value adjusted with FDR, α=0.05). B) Overlap between significantly over-targeted gene sets. C) Most targeted genes in MSCs and HSPC. IS enrichment scores highlight each calculated dataset’s top 10 most targeted genes. Genes associated with documented clonal expansion in gene therapy clinical trials are included in the left part of the heatmap, marked by a red line under the gene name. The color in the heatmap represents the IS enrichment, quantified as the log odds (regression coefficient estimate) associated with the gene effect. D) Zoom in into over-targeted genes: Detailed view of two over-targeted genes reveals cell type-specific and shared integration patterns: *NPLOC4*: Shared over-targeted gene among all cell types. *PLEC*: Over-targeted exclusively in both BM MSC and Ad MSC.

In BM MSC, we found 5313 genes had at least 1 IS within their extended gene body, of which 1134 were significantly over-targeted. In Ad MSC, 718 genes out of 5401 were significantly over-targeted.

As shown in Figure 4B, 157 genes were over-targeted in all cell types, 109 genes appear to be MSC specific, and the overlap between BM MSC and HSPC (295 genes) is higher than the shared genes between Ad MSC and HSPC (134 genes).

We defined a set of High-Risk genes that includes cancer-associated genes and genes related to IM events and verified how their targeting rates varied among cell types (Sadelain et al., 2011). The heatmap in Figure 4C highlights how the IS patterns are very similar, especially on the highly targeted High-Risk genes. The one exception was *SPPL2B* which was specific for BM MSC. Interestingly, genes associated with documented clonal expansion in humans, such as *LMO2*, have no detected IS, and genes such as *MECOM* and *HMGA2* had scores higher in HPSC than MSCs.

We provide a more detailed representation of the IS distribution around two genes in Figure 4D. *NPLOC4* appeared among the top ten targeted genes across all datasets, while *PLEC* is highly enriched in IS in both BM MSC and Ad MSC datasets but there was no IS noted in the HSPC dataset. The complete results of the analysis are available in the Suppl. Files, the genome-wide gene targeting score visualization is provided in Suppl. Figure 3 and the pathway analysis for each dataset in Suppl. Figure 4-6.

### Differential gene targeting analysis highlights differences between MSC and HSPC and minor differences within MSC integromes

In addition to the gene sets intersection analysis, MELISSA allows us to compare the safety profiles in two cell types with a dedicated model, avoiding loss of relevant information and focusing exclusively on detecting the differences between, for example, a novel and a reference integrome.

First, we compared BM MSC to HSPC (Figure 5A) and identified 119 genes over-targeted in BM MSC and 47 in HSPC. The most over-targeted gene in BM MSC was *PLEC* (Figure 4D), while *DACH1* was the most over-targeted gene in HSPC and showed no IS in BM MSC. The waterfall plot in Figure 5B further investigates the differential targeting of the High-Risk gene set in BM MSC and HSPC. Despite 29 high-risk genes being over-targeted in BM MSC (top genes: *SPPL2B*, *PPARD*, *STAG2*, *SETD2*) and 13 in HSPC (top genes: *GRB2*, *PTPRD*, *IKZF2*), the risk profiles of the two integromes are well-balanced overall, with an average risk score equal to 0.4 (0=equal risk). Pathway analysis calculated using gene targeting scores is reported in Suppl. Figure 7.

**Figure 5:**
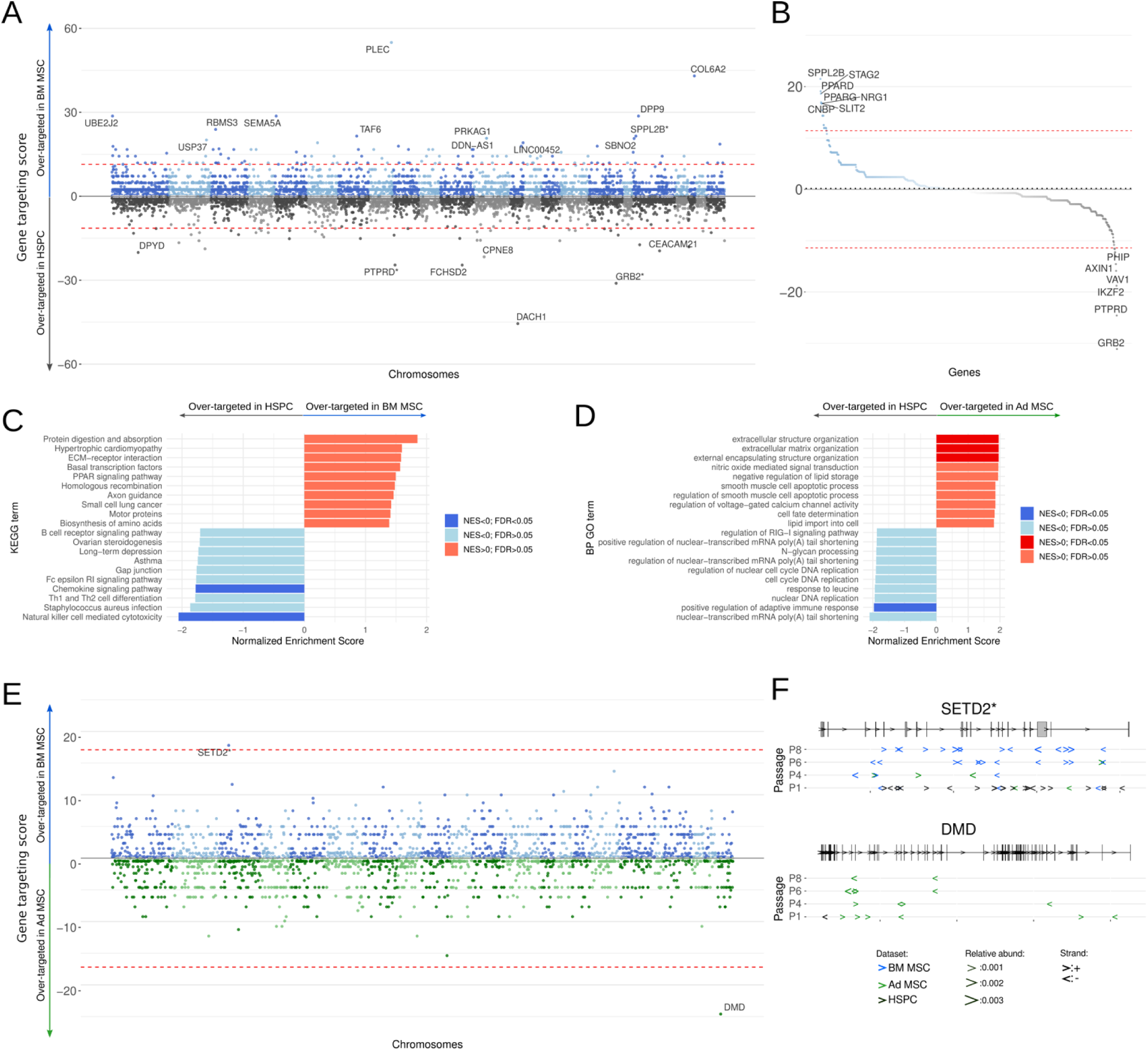
Differential gene targeting analysis. A) BM-MSC/HSPC analysis: Differential gene targeting analysis comparing BM-MSC and HSPC. Each dot represents a gene. If the gene is enriched in the integration site in BM-MSC (HSPC), it is plotted in the blue (grey) section. Genes marked with a * are included in the high-risk gene list. The dashed red line represents the threshold for statistically significant enrichment (LRT p-value adjusted with FDR, α=0.05). B) BM MSC/HSPC waterfall plot for high-risk genes. Integration site enrichment scores for genes included in the high-risk genes are ranked by targeting score. The dashed red line represents the threshold for statistically significant enrichment (LRT p-value adjusted with FDR, α=0.05). C) BM MSC/HSPC KEGG pathway enrichment analysis. Pathway analysis was calculated using differential gene targeting scores. D) Ad MSC/HSPC Gene Ontology enrichment analysis. GO pathway analysis was calculated using differential gene targeting scores. E) Ad MSC/HSPC analysis. Differential gene targeting analysis comparing BM MSC and Ad MSC. Each dot represents a gene. If the gene is enriched in the integration site in BM MSC (Ad MSC), it is plotted in the blue (green) section. Genes marked with a * are included in the high-risk gene list. The dashed red line represents the threshold for statistically significant enrichment (LRT p-value adjusted with FDR, α=0.05). F) Zoom in into differentially over-targeted genes. A detailed view of two over-targeted genes reveals tissue-specific gene targeting: *SETD2* over-targeted specifically in BM MSC. *DMD* is over-targeted, specifically in Ad MSC.

When evaluating genes with high IS rates in the Ad MSC and HSPC analysis (Suppl. Figure 8-9), 9 out of 51 genes in the Ad MSC group and 5 out of 21 in the HSPC group were present in the High-Risk gene list.

The KEGG and the GO analysis in Figure 5C,D were calculated using the gene’s IS rates scores from the comparisons between MSCs and HSPC. The analysis shows that IS rates can discriminate cell type-specific pathways, such as in immune responses and signaling pathways in HSPC and extracellular matrix regulation in Ad MSC.

When comparing BM MSC to Ad MSC (Figure 5E, Suppl. Figure 10), we found only 2 differentially targeted genes. *SETD2* (Figure 5F), a critical chromatin regulator with potential tumor suppressor activity (Chen et al., 2020) is over-targeted in BM MSC. The *DMD* gene was over-targeted in Ad MSC. Despite the *DMD* gene not being part of the High-Risk gene sets, it is noteworthy that a growing body of evidence suggests a potential role of the *DMD* gene in tumor development and progression (Alnassar et al., 2023; Naidoo et al., 2022).

### Different gene sets influence MSC clone growth dynamics in-vitro

While analyzing gene-targeting rates reveals potential safety risks, it does not directly assess whether IS within specific genes are likely to alter cell behavior. To address this crucial safety concern, we can quantify how targeting a particular gene impacts the fitness of individual cell clones. MELISSA’s model tracks the growth of clones with IS within specific genes, comparing them to average growth rates and potentially across different conditions. This approach helps discern whether clone selection is driven by stochastic factors or a direct consequence of the IS.

Using a gene/growth association score calculated from the single condition statistical analysis, we measured the impact for genes targeted in BM MSC and Ad MSC. Interestingly, the gene scores shown in Figure 6A reveal distinct sets of top genes associated with growth for each cell type. In contrast, genes with documented potential for clonal expansion have no significant impact. In the Ad MSC dataset, we identified more genes significantly affecting clone growth rates (Ad MSC: 48, BM MSC: 32) and a higher number of High-Risk genes (Ad MSC: 17, BM MSC: 7).

**Figure 6:**
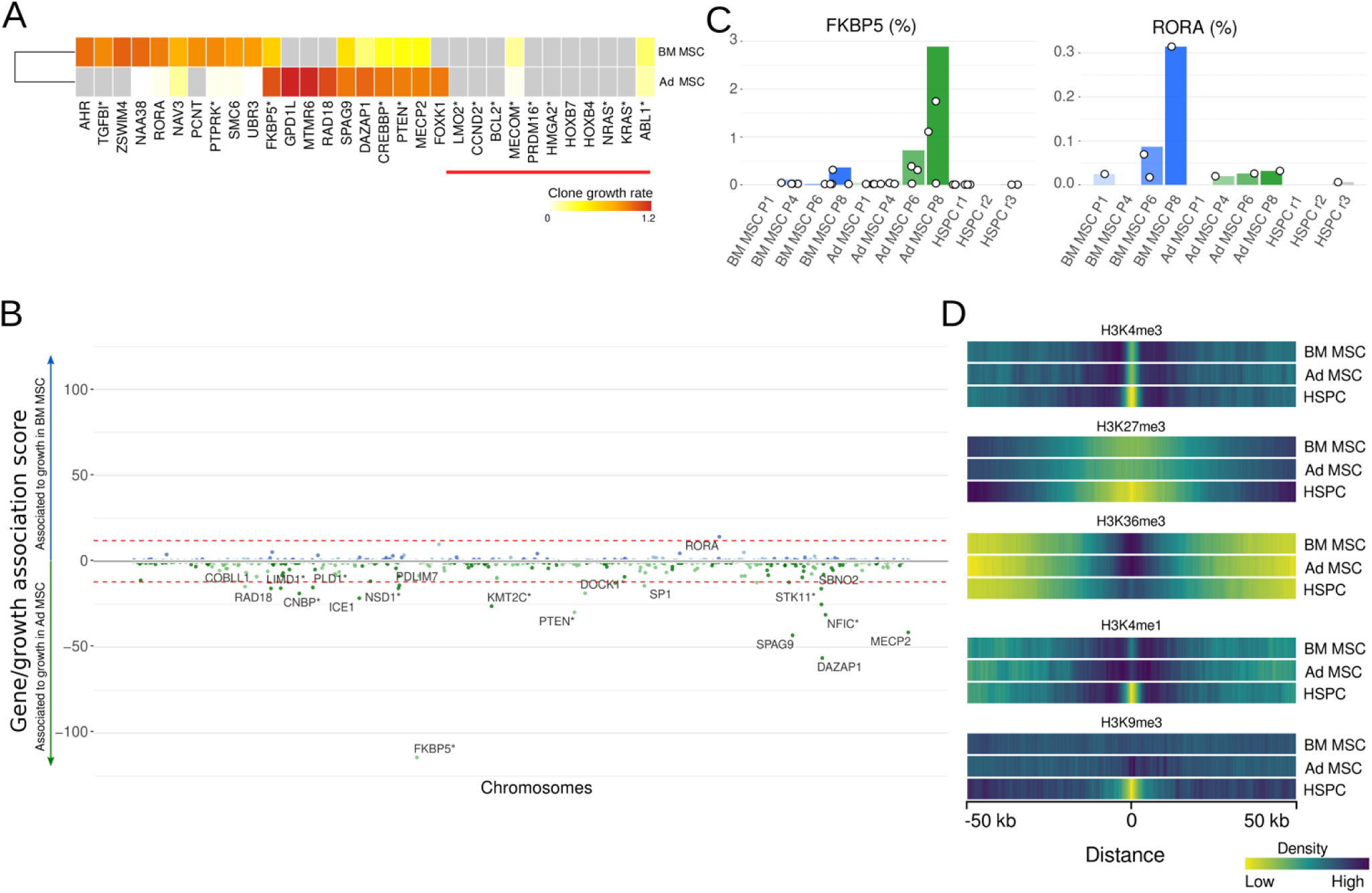
Integration site gene targeting effects on clone growth dynamics. A) Targeted genes with the highest positive impact of clone growth. Clone growth rate scores for the top 10 genes identified using the clone growth dynamics modeling in a single group setting. Genes associated with documented clonal expansion in gene therapy clinical trials are included in the left part of the heatmap, marked by a red line under the gene name. The color in the heatmap represents the estimated clone growth rates, measuring the average increase in clone size at each passage for IS mapping to a specific gene. B) Enrichment score for differential clone growth dynamics in BM MSC and Ad MSC. The plot illustrates the growth association score, quantifying the differential impact of integration site targeting genes on the growth dynamics of clones within BM MSC and Ad MSC samples longitudinally over passages. Each dot represents a gene. If IS in the gene increases clone growth in BM MSC (Ad MSC), it is plotted in the blue (green) section. Genes marked with a * are included in the high-risk gene list. The dashed red line represents the threshold for statistically significant differential clone dynamics (LRT p-value adjusted with FDR, α=0.05). C) Zoom in into genes with the most significant growth association score. Detailed view of the relative contribution during the in-vitro culture of IS within *FKBP5* (Ad MSC) and *RORA* (BM MSC). The bar graph represents the IS’s cumulative contribution (%) within the gene, while dots correspond to individual clones. D) Integration site association with histone modifications in the different cell types. Comparison of the distance between IS datasets and the (top to bottom) H3K4me3, H3K27me3, H3K36me3, H3K4me1, and H3K9me3 histone modification profiles in the corresponding cell types. The histogram (bin size 300bps) of distances is generated and converted into a heatmap.

Figure 6B highlights a key difference: IS in 19 genes show a significant positive impact on Ad MSC clone growth, with *FKBP5* showing the most substantial effect. Conversely, only one gene, *RORA*, positively influenced BM MSC growth. Figure 6C further details these findings, visualizing the individual and cumulative contributions of clones harboring IS events within FKBP5, reaching a 3% total contribution with 9 IS, while the *RORA* effect is less pronounced and limited to 0.3%.

### Integration preferences in MSC and HSPC show different correlations with epigenetic marks

MELISSA offers several tools to help in the biological interpretation of the findings derived from the IS rates and clone fitness analysis.

We investigated the relationship between the distribution of IS and chromatin status in a cell-type-specific manner by taking advantage of the Roadmap Epigenomics project, which published Chromatin ImmunoPrecipitation followed by sequencing (ChIP-seq) data on the histone modifications H3K4me1, H3K4me3, H3K9me3, H3K27me3, H3K36me3. We utilized data reports for BM MSC, Ad MSC and mobilized CD34+ HSPC (Suppl. Table 4). Our analysis assessed the enrichment of these epigenetic marks within a 100 kb window centered on the IS, similar to the analysis proposed in (Yan et al., 2023). A heatmap representation of distance histograms is shown in Figure 6D.

The results for the HSPC dataset are consistent with the profiles previously reported in the literature for all the six marks analyzed. For the open chromatin markers H3K4me3 and H3K36me3 and the repressive mark H3K27me3, we found a consistent pattern of distance distribution in both MSC cell types and HSPC, with the typical ± 2 kb sharp short-distances depletion for H3K4me3 and the progressive and smooth short-distances enrichment and depletion for H3K36me3 and H3K27me3 respectively. However, for the marker of active enhancers H3K4me1 and the repressive mark H3K9me3, MSC cell types and HSPC IS showed different patterns. In HSPC, short distances (±2kb) were underrepresented for both histone modifications, yet this pattern was not found in BM MSC and Ad MSC.

### Gene targeting and clone fitness analysis in Gene Therapy Clinical Trials

For each patient, IS tables were constructed from myeloid (Monocytes and Granulocytes) and lymphoid (B, T, and NK cells) subpopulations isolated from peripheral blood samples collected at least twice post-therapy. Figure 7A summarizes the data, presenting the number of IS and the cumulative clone size for each cell type, time point, and patient. The datasets exhibit considerable heterogeneity in size across trials and patients, with the β-thal patient showing a significantly higher number of IS compared to SCD, and within the WAS trial, the WAS2 patient displaying a higher IS count than the others.

**Figure 7.**
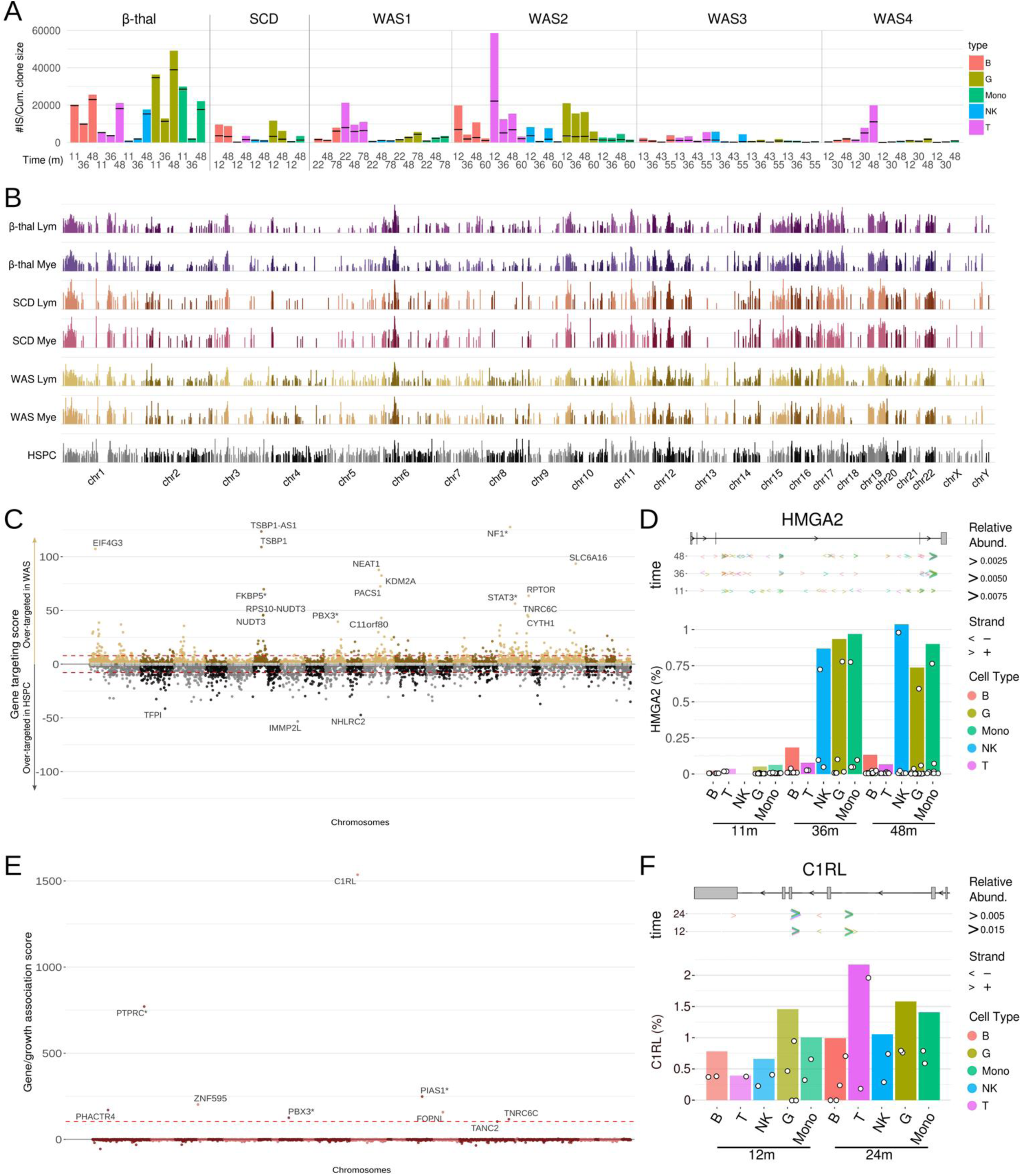
Gene targeting and clone fitness analysis on gene therapy clinical trials IS data. A) Datasets summary. Number of IS (black horizontal lines) and cumulative clone sizes (bar) for each sample (trial, patient, cell type and time point). B) IS enrichment scores for genes found significantly over-targeted (LRT p-value adjusted with FDR, α=0.05) in at least one data set using the single-group gene targeting analysis. Each bar corresponds to a gene. Chromosome location is reported on the x-axis. C) Differential gene targeting analysis comparing WAS and HSPC. Each dot represents a gene. If the gene is enriched in the integration site in WAS (HSPC), it is plotted in the brown (grey) section. Genes marked with a * are included in the high-risk gene list. The dashed red line represents the threshold for statistically significant enrichment (LRT p-value adjusted with FDR, α=0.05). D) Focus plots for clones harboring IS in the *HGMA2* gene in the β-thal trial. Top: IS position. Bottom: Individual and cumulative abundance. E) Gene/growth association in the SCD trial. The gene/growth association score provides a metric for assessing the significance of the fitness advantage or disadvantage conferred by IS within a gene. Genes marked with a * are included in the high-risk gene list. The dashed red line represents the threshold for statistically significant differential clone dynamics (LRT p-value adjusted with FDR, α=0.05). F) Focus plots for clones harboring IS in the *C1RL* gene in the SCD trial. Top: IS position. Bottom: Individual and cumulative relative abundance.

MELISSA models, which utilize logistic regression, effectively account for these data imbalances and the effects of progressive clone selection over time. Figure 7B displays the estimated targeting rates for genes significantly over-targeted in at least one analysis. Miami plots for individual clinical trials are provided in Suppl. Figure 11.

Compared to the 1,944 genes significantly over-targeted in HSPC, fewer over-targeted genes were identified in the clinical trial datasets. No major differences were observed between myeloid (β-thal: 835, SCD: 687, WAS: 1,215) and lymphoid lineages (β-thal: 875, SCD: 702, WAS: 1,285). However, the clinical data exhibited a broader range of gene targeting scores than HSPC. Notably, there was a strong increase in highly targeted genes in specific genomic regions, such as chromosome 6, and a consistent decrease in targeting scores in regions like chromosomes 4 and 18 across lineages and trials.

To assess whether gene targeting rates differ significantly between clinical samples and control HSPC data, we employed MELISSA differential models. Figure 7C highlights the comparison between WAS, with data from four patients and the longest follow-up period (Suppl. Figure 12 for β-thal and SCD), to determine whether IS within specific genes are more likely to pass the bottleneck of engraftment, characterizing the integrome of long-term repopulating HSCs. In alignment with the single-group analyses, several broad genomic regions were significantly differentially targeted. For instance, regions such as chr6 p22.1-p21.31, chr1 p36.13-p36.11, and chr11 q12.2-q13.4, while enriched in targeted genes in HSPC, exhibited such an increase in their relative contribution in the clinical datasets that clones harboring IS in these regions have a significantly higher probability of long-term engraftment. Conversely, genes in chr13 and chr18, significantly over-targeted in HSPC, are likely lost upon in vivo selection.

We conducted a clone fitness analysis across all available clinical trial data, incorporating cell type as a covariate to identify evidence of IM. Given the limited number of patients in the clinical data, to minimize potential false-positive gene associations caused by stochastic clonal selection (without mechanistic interaction), we implemented an additional robust analysis step that filters out gene/clone growth associations that are not significant after excluding measurements from the most abundant clone in the tested gene. Further details on this procedure are provided in the Methods section.

In the β-thal dataset, IS in three genes were identified as potentially altering clone fitness and conferring a proliferative advantage (Figure 7D, Suppl. Figure 13). The most significant, *HMGA2* (LRT-score: 2,732, adjusted p-value < 1e-16), encodes the AT hook 2 transcription factor, known for binding AT-rich DNA regions. This IM event, extensively described in previous studies (Cavazzana-Calvo et al., 2010; Kumar et al., 2019; Mansoori et al., 2021), suggests that IS in the last intron generates mRNA lacking let-7 miRNA binding sites, leading to reduced *HMGA2* degradation and driving proliferative advantage. Despite the expansion of *HMGA2*-integrated clones, no progression to a leukemic phenotype was observed, and the event was classified as benign due to its stabilization over time. MELISSA’s statistical model detected this IM event using IS tables from just three time points (11, 36, and 48 months), with a cumulative IS contribution in *HMGA2* of only 1%. As shown in Suppl. Figure 14, this effect is specific to the myeloid lineage, consistent with published data. The second most significant gene, *SFPQ* (LRT-score: 458, adjusted p-value < 1e-5), is an epigenetic regulator with histone deacetylase binding activity.

In the SCD dataset (Figure 7E), six genes were identified with significant increases in growth rate. The most significant, *C1RL* (LRT-score: 1,558, adjusted p-value < 1e-11), encodes Complement Component 1 Receptor-Like protein, involved in immune response modulation. A detailed representation of IS location and abundance in *C1RL* is shown in Figure 7F. Despite a maximum cumulative contribution of 2%, the consistent increase across multiple independent clones and lineages suggests a potential mechanism linking *C1RL* to IM events in gene therapy. Additionally, *C1RL* has been significantly associated with survival in a cohort of acute myeloid leukemia patients and is widely expressed in myeloid leukemia cell lines, including K-562 (chronic myelogenous leukemia), THP-1 (myelomonocytic leukemia), HL-60, and NB4 (acute promyelocytic leukemia), as well as glioma (Laverdière et al., 2016; Wang et al., 2020).

## Discussion

Ensuring the safety of gene and cell therapies is a top concern for all stakeholders in the field, from researchers and regulators to patients and their advocates. While IS analysis is a cornerstone of evaluating safety outcomes in preclinical and clinical studies using cell products transduced with viral vectors, the existing analytical frameworks primarily focus on sample report generation (Sherman et al., 2016), data handling (Merelli et al., 2023), and modeling of clonal tracking data (Espinoza et al., 2021; Pais et al., 2023; Wagner et al., 2022). This leaves a critical gap in evaluating safety concerns such as gene targeting rates and the presence of clones with altered fitness and proliferative advantage. To address this gap, we developed a novel analytical framework named MELISSA.

MELISSA is designed as an R package and facilitates the analysis, combination, and comparison of IS datasets generated using diverse conditions and complex experimental design. This powerful tool allows researchers to leverage the flexibility of regression models, generating gene-referenced quantitative scores that measure both targeting rates and their effect on clone fitness, and include dedicated functionality to explore the biological relevance of the results.

The simulation study presented in Figure 2 highlights the remarkable performance of our methodology. Across all scenarios, the PPV increased as both effect size and sample size grew, showing the consistency of the testing procedure. This is true for target genes of all simulated lengths, although, for long *TestGenes* (40-80 Kb), the PPV shows a stronger increase with effect size.

The PPV curves for gene targeting (Figure 2A,B) are similar, despite the differential analysis using twice the data of the single-group analysis. In our simulations, genes with fewer than 2 IS were excluded (users defined parameters). Shorter genes (5–10 Kb) were less likely to be included in the single-group analysis but, when tested, were more easily detected as significant. For genes of typical length (20–40 Kb), effect sizes of 4 (single-group) and 2 (differential) ensured sufficient coverage, though detection rates exceeded 50% only at effect sizes of 16.

In terms of clone fitness analysis, which is crucial for detecting risk of IM, our modeling demonstrated the best performance across all metrics in the single-group scenario. It excelled in terms of PPV, coverage, and detection rate, even for shorter *TestGenes* and low effect sizes, highlighting its sensitivity. In differential settings, the detection rate was lower due to the model leveraging in particular the growth rates of a restricted number of clones—those located in *TestGenes* in the two groups.

Recent research highlights the potential of MSCs for gene therapy applications. Their immunomodulatory and regenerative properties, ease of isolation, rapid ex vivo expansion, and multilineage differentiation capabilities make them promising vehicles for targeted therapies in various contexts, including cancer gene therapy (Mohammadi et al., 2016). However, studies have revealed heterogeneity among MSCs derived from different tissues, suggesting this variation might contribute to the diverse levels of efficacy and outcomes observed in animal studies and MSC-based clinical trials.

To address this limitation, we characterized and make available to the scientific community the first detailed LV integrome in BM MSCs and Ad MSCs. We then compared their safety profiles, using HSPC transduced with the same LV vector as a reference. We chose HSPCs due to the extensive data available on their LV integration distribution and associated effects, providing a well-established benchmark for comparison.

Figure 3 highlights critical differences in clonal dynamics between the cell types despite both BM MSCs and Ad MSCs demonstrating high transduction efficiency. BM MSCs exhibited a more stable clonal composition, suggesting a more homogeneous cell population that better adapts to the culture conditions. Ad MSCs displayed a marked decline in diversity and IS count over the culture period, suggesting a progressive loss of transduced cells and a limited proliferative activity compared to BM MSCs. Therefore, we did not identify an IS profile that could discriminate more primitive culture-initiating cells from the whole MSC cell population. This is in contrast to the study by (Selich et al., 2016) where primitive culture-initiating cell expansion was observed. It must be noted that MSC in their study were obtained from whole umbilical cord pieces and may well have a different phenotype from both BM and Ad MSCs.

Encouragingly, the distribution of IS with respect to gene annotation was consistent across cell types. We found that the high propensity of LV to integrate within gene bodies extends to MSC and this feature strengthen the relevance of our gene-based approach, with only 15% of the IS data points not considered in our analysis due to their intergenic position.

Our study generated a valuable resource for researchers working with LV-HSPC gene therapies consisting of a dataset composed of over 35,000 IS across three replicates. This dataset closely aligns with existing knowledge on HSPC integromes regarding RIG and overall association with the epigenetic profile (Figures 4, 6D).

Analysis of gene-specific IS rates revealed both shared and MSC-specific integration preferences. Among the shared targeted genes, *NPLOC4* serves as an example, showing high targeting not only in MSCs but also in ex vivo transduced HSPCs from healthy donors and clinical trial subjects (Aiuti et al., 2013; Biffi et al., 2013; Cartier et al., 2009; Hacein-Bey Abina et al., 2015), as well as in LV transduced T cells (Malach et al., 2023). In contrast, *PLEC* exemplifies an MSC-specific target, as it was the most frequently targeted gene in MSCs and absent in the IS profiles of HSPCs. The frequent targeting of *PLEC* in MSCs is particularly noteworthy given previous findings of high-frequency mutations in the *PLEC* gene in bone marrow-derived MSCs from patients with Acute Myeloid Leukemia, suggesting a potential link between *PLEC* dysfunction and MSC involvement in AML development (Fracchiolla et al., 2017). This raises potential safety concerns for LV-modified MSCs, especially regarding PLEC-targeted integrations.

Furthermore, our study revealed similar targeting of high-risk cancer genes across MSC and HSPC cell types with some distinctions. For example, *SPPL2B*, a protease that cleaves the transmembrane domain of tumor necrosis factor-alpha to release the intracellular domain and is known for its role in immune regulation and inflammation, was targeted exclusively in BM MSC. More deep studies involving in vivo experiments will be crucial to identify MSC-specific IM risk genes, as our results found low targeting rates and no clonal expansion associated with genes previously known to cause clonal expansion in animal models and human subjects involved with LV/RV HSPC gene therapy.

Gene therapy safety assessment often relies on comparative IS data analysis from different sources. This can involve, as demonstrated in this study, comparing gene targeting rates between alternative cell sources (e.g., BM MSC vs. Ad MSC), vector designs, or transduction protocols.

Figure 5 highlights MELISSA’s capability of capturing differentially targeted genes. This allows for in-depth investigation of individual genes and calculation of genome-wide high-risk scores. Since LV integration often correlates with gene expression and active enhancers, comparing MSCs and HSPCs revealed differentially targeted genes potentially involved in cell type-specific pathways. In the comparative analysis focusing on MSC cells, only *SETD2*, an epigenetic regulator with potential tumor suppressor activity (Chen et al., 2020), was identified as differentially targeted in BM MSC, and *DMD* in Ad MSC. However, as demonstrated in the simulation study, the ability of MELISSA to detect differentially targeted genes is contingent upon the strength of the effect and the size of the dataset. It is plausible that by augmenting the number of IS retrieved from the MSC experiment, more differentially targeted genes with weaker associations could be identified, suggesting the potential for further exploration for a more conclusive safety assessment comparison.

Using MELISSA’s clone fitness analysis, we explored the relationship between LV integration and clone fitness in MSCs. Figure 6 demonstrates that distinct sets of genes significantly impacted in-vitro clone growth dynamics within each MSC population. Notably, genes previously linked to clonal dominance or IM in HSPC-based gene therapy exhibited minimal influence on MSC expansion. This reinforces the crucial need for cell type-specific considerations when evaluating gene sets for their potential impact on clone fitness and associated safety risks.

The differential analysis identified a single gene, *RORA*, significantly impacting clone growth in BM MSCs. Ad MSCs, however, displayed a more complex picture with more significant genes and stronger effects. Precisely, clones with IS in *FKBP5*, a known oncogene, are estimated to double their contribution with each passage of in-vitro culture. However, it is important to note that these associations might also be influenced by the observed consistent and significant loss of transduced cells in Ad MSCs. This reduces overall cumulative clone size and exacerbates the contribution of surviving clones.

This is the first report of insertion site analysis of MSC. While the differences noted in BM MSC and Ad MSC are interesting, further studies evaluating different donors are warranted. Gene modification of MSC is likely to transfer chemokine, cytokine, or other genes aimed at improving their potency but could also impact fitness and therefore worthy of future study. It is also worth noting that LV vectors are currently the most prevalent delivery vehicle for patients treated with FDA approved gene therapies and in many ongoing clinical trials.

The association observed between histone modifications and IS distribution in HSPCs aligns with findings reported in SCID-X1 (Yan et al., 2023) and wild-type HIV (Lelek et al., 2015). However, the epigenomic profiles of MSCs only exhibit partial correlation with their respective integromes. Notably, H3K4me1 and H3K9me3 display distinct patterns of association between MSCs and HSPCs with their respective integromes. This inconsistency could be explained by the co-localization of H3K9me3 and H3K4me1 specific to HSPCs or the availability of tethering factors differing between cell types. Further studies are needed to fully understand the biological reasons behind this lack of consistency. However, it highlights the importance of in-depth characterization of how host cell status drives the IS selection process in a cell type-specific manner.

Our analysis across multiple gene therapy clinical trials, as shown in Figure 7, reveals significant heterogeneity between the integrome observed in vitro within HSPCs and that in myeloid and lymphoid subpopulations isolated from the peripheral blood of GT patients. These differences can largely be attributed to the nature of the cell populations and the timing of data collection. HSPCs are a diverse group, comprising both short-term progenitors and long-term repopulating HSCs. The HSPC data tables were generated shortly after transduction, without undergoing the selective pressures of engraftment seen in clinical data. Conversely, the clinical IS data, generated from peripheral blood samples at least 11 months post-gene therapy, are more reflective of the integrome of long-term HSCs. The consistent patterns of clonal selection observed across different trials, particularly in spatial distribution, suggest that these changes are driven by in vivo clonal selection processes rather than by the specific disease context.

The identification of known gene/growth associations, such as *HMGA2* in the β-thal trial, provides evidence that the MELISSA model can capture mechanistic effects despite the limited data available. In the SCD trial, we found clones with IS in *C1RL* to exhibit a similar expansion rate to that of *HMGA2* in β-thal. This highlights the importance of considering significant associations, especially in situations where data is limited to a single or few patients, as potential indicators of IM events that warrant close monitoring to differentiate between benign and malignant proliferative events and avoid false-positive associations. However, we anticipate that as the number of patients in a dataset increases, MELISSA will be able to detect more subtle effects and further reduce the incidence of false positives, enhancing these findings’ robustness.

While MELISSA offers valuable insights, it is important to acknowledge its limitations. Firstly, the current gene-centric approach may overlook potential safety concerns arising from IS within non-coding genome regions. We are actively developing methods to address this limitation and provide a more comprehensive, gene-location independent, analysis. Additionally, different viral vectors may exhibit varying preferences for specific genomic elements, potentially resulting in a higher proportion of uncharacterized integration sites. In such cases, the genomic intervals to be tested can be extended to include alternative genomic annotations, such as chromatin accessibility profiles or patterns of histone methylations (gene regulatory regions, enhancers).

In conclusion, we present MELISSA, a novel framework for analyzing IS data to assess the safety of viral vector-mediated gene and cell therapies. MELISSA offers diverse functionalities for gene targeting analysis, differential analysis, clone growth analysis, and visualization. Applying MELISSA MSCs revealed distinct clonal dynamics, cell type-specific gene targeting, and the importance of cell type considerations for evaluating safety. We envision MELISSA to become a valuable tool for empowering researchers, clinicians, and regulators to ensure the safety of novel gene and cell therapies.

## MATERIALS AND METHODS

### Lentiviral vectors

A third-generation^12^ LV encoding EGFP (pcDNA-CSCGW^43^ kindly provided by P. Zoltick, Children’s Hospital of Philadelphia, Philadelphia, PA) was produced in a 5 L shaking flask (Thompson, Oceanside, CA, USA) using the packaging plasmids pMDL,^12^ pRSV-Rev,^12^ and pMD.G^44^. Serum-free media adapted suspension HEK293T(IUS) cells, derived from HEK293T cells (Stanford University), were cultured in Expi293 media (Gibco, New York, USA) at 2.5 x10^5^ cells/mL using shake flasks in a 37 degree C 5% CO_2_ incubator for at least 2 passages before transfection. Cells were numerated on the day of transfection and resuspended in TransFX media (Cytiva, Marlborough, MA, USA) supplemented with 4 mM of GlutaMAX (Gibco, New York, USA). The transfectant, containing OptiMEM (Gibco, New York, USA), 4 plasmid DNA, and PEIpro (Polyplus, Illkirch-Graffenstaden, France), was prepared at 5% of the total cell culture volume. Per 10^6^ cells, 2 µg of total plasmid DNA were added. Plasmids pcDNA-CSCGW, pMDL, pMD.G, and pRSV-Rev were mixed into OptiMEM at the weight ratio of 4:2:1.3:1, respectively. PEIpro was added to the plasmid mix at a ratio of 1 µL of PEIpro per 1 µg of total plasmid DNA and allowed to polyplex for 15 minutes at room temperature. The transfectants were added to the HEK293T(IUS) cells for a final concentration of 3 x 10^6^ cells/mL; transfected HEK293T(IUS) cells were cultured in a 37°C incubator supplied with 5% CO_2_ for 16 hours. After 16 hours of incubation, TransFX media containing 10 mM of sodium butyrate (Cayman Chemical Co., Ann Arbor, MI, USA) was added to the cell culture at a ratio of 1:1. Forty-eight hours post-transfection, culture media were harvested and filtered through 0.4 µm filters to remove cell debris. Filtered LV was allotted into cryovials, 1 mL/vial, and stored at –80°C freezer until use.

### Transduction of MSC

Bone marrow-derived MSCs (BM-MSCs) were acquired from ATCC and grown in Rooster Media (Rooster Bio). Adipose-derived MSCs (AT-MSCs) were acquired from ATCC and grown in DMEM/F12(Gibco) supplemented with 5% FBS(Cytiva), 10ng/mL FGF-2 (R&D Systems), 5ng/mL EGF (R&D Systems), and 250uM ascorbic acid. The cells were maintained in a 37°C 5% CO_2_ incubator. For both BM and ASC experiments, cells at P1 were seeded at 10,000 cells per cm^2^ in 6-well dishes (Corning) for a control and transduced group. The following day, the media was aspirated from the transduced group and 1mL of LV diluted 1:100 was added to the wells along with Polybrene (final concentration 8 ug/mL). Four hours later, the LV solution was aspirated, and fresh culture media was added. Once the wells reached 70-80% confluence, the cells were washed with DPBS(Gibco), harvested using TrypLE(Gibco), and seeded between 3,000 and 5,000 cells per cm^2^. At passage 4, the vector copy number (VCN) per cell for BM MSC and Ad MSC was 3.91 and 2.69, respectively. At passage 8, the VCN per cell was 3.45 and 2.82 for BM MSC and Ad MSC, respectively. This procedure was repeated for both the control and transduced samples until P5. At P5, some of the control wells were transduced in the same manner as previously described. The passaging procedure was continued for all three groups until P11. The cells were collected at various time points for insertion site analysis. The cultures were also observed using a fluorescent microscope to observe GFP expression.

### Transduction of CD34+ HSPCs

GCSF-mobilized peripheral blood-derived (mPB) CD34+ HSPCs were pre-stimulated for 24 hours and two consecutive overnight transduction were performed at a concentration of 5 x 10^5^ cells/mL at a multiplicity of infection (MOI) of 75 per transduction in the presence of the transduction enhancers synperonic F108 (200 µg/mL final concentration, Sigma) and protamine sulfate (4 µg/mL final concentration, Sigma) in serum-free stem cell growth medium (CellGenix® GMP SCGM) supplemented with 1% Penicillin/Streptomycin (Gibco), 1% L-glutamine (Gibco), 100 ng/mL rhSCF, 100 ng/mL rhTPO and 100 ng/mL rhFlt3L (all cytokines: Peprotech). The cells were cultured in supplemented SCGM for ten days before genomic DNA isolation.

### Droplet digital PCR for vector copy numbers in transduced cells

Sample DNA was extracted from cells using QIAamp DNA Mini Kit (Qiagen). The primers and probe nucleotides sequences for detecting LV are forward primer LV-F (5’-ACTTGAAAGCGAAAGGGAAAC-3’), reverse primer LV-R (5’-CACCCATCTCTCTCCTTCTAGCC-3’), and probe TP-LV (5’-6FAM-AGCTCTCTCGACGCAGGACTCGGC-TAMRA-3’). Apolipoprotein B (APB) gene was used as an endogenous control; the primers and probe sequences for APB are forward primer APB-F (5’-TGAAGGTGGAGGACATTCCTCTA-3’), reverse primer APB-R (5’-CTGGAATTGCGATTTCTGGTAA-3’), and probe TP-APB (5’-VIC-CGAGAATCACCCTGCCAGACTTCCGT-TAMRA-3’). The primers and probe nucleotides were synthesized by a commercial vendor (Integrated DNA Technologies).

Droplet digital PCR (ddPCR) was performed on the Bio-Rad QX200 system to quantify vector copy numbers in transduced MSCs by following the manufacturer’s procedures. The controls are no-template controls (water), negative controls (DNA from untransduced CD34(-) cells), and positive controls (150000 copies of target sequences). Each sample was tested in triplicates. An excess master mix of reagents without DNA was prepared according to the total number of testing articles (including controls) and the following conditions – for each reaction, a final concentration of 1x no dUTP ddPCR super mix for probe (Bio-Rad), 900 nM of LV-F, 900 nM of LV-R, 250 nM of TP-LV, 900 nM of APB-F, 900 nM of APB-R, 250 nM of TP-APB, and 5 units (in 1 µL) of HindIII-HF restriction enzyme (New England BioLabs) were assembled. The master mix was distributed to a microtube before adding 1 µg of testing article DNA; DNase/RNase-free water was added to bring the final volume to 20 µL in each reaction. The 20 µL samples were loaded into a cartridge with 70 µL of generation oil loaded in all the bottom wells for droplet generation in a droplet generator (Bio-Rad). The formed droplets were transferred to a 96-well ddPCR plate (40 µL droplets/well) before being sealed with a pierceable foil heat seal at 180°C using a heat sealer (Bio-Rad). The sealed plate was placed on a C1000 Touch Thermal Cycler (Bio-Rad), and the PCR was run using the following program – Step 1 (95°C for 10 minutes), Step 2 (40 cycles of 94°C for 30 seconds followed by 60°C for 1 minute), and Step 3 (98°C for 10 minutes). After Step 3, the thermocycler holds at 12°C until reading the droplets using a QX200 droplet reader (Bio-Rad). The data was analyzed using QX200 Manager Software (Bio-Rad).

### DNA Library preparation for integration site analysis

DNA library for integration site analysis was prepared according to the INSPIIRED^31^ protocol unless otherwise noted here. The genomic DNA from LV-transduced cells was extracted and eluted in DNase-/RNase-free water with QIAamp DNA Mini Kit (Qiagen) by following the manufacturer’s procedure. The extracted genomic DNA was fragmented at the Indiana University School of Medicine Center for Medical Genomics Service Core (IUSM-CMG). DNA fragmentation was performed by sonication using the Covaris ME220 sonication system with 40 Watts peak power, 10% duty factor, 1000 cycles/burst at 25 degree C for 30 seconds, Oligonucleotides (linkers, primers, and blocking oligos) for DNA library preparation were synthesized, according to the INSPIIRED protocol,^31^ by Integrated DNA Technologies, Inc. The DNA library was evaluated on a Bioanalyzer High Sensitivity DNA Analysis at IUSM-CMG to ensure library quality. A maximum of 5 prepared DNA libraries were sequenced using MiSeq Reagent Kit v2 300-cycle (Illumina) on the MiSeq system at IUSM-CMG.

### Statistical modeling

The four statistical analyses considered in this manuscript are the following.

A) Gene targeting. The tested gene has a higher IS rate than the rest of the genome. B) Differential gene targeting. The rate of IS in the tested gene is different in the two groups. C) Clone fitness. In the tested gene, clone sizes grow faster (on average, over time) than in the rest of the genome. D) Differential clone fitness. In the tested gene, there is a difference between group 1 and 2 in clones’ growth rate. For analyses A) and B), the presence/absence of an IS at each genomic coordinate is modeled as a binary variable (1 if an IS maps to the specific location and 0 otherwise). This binary variable is then considered as the response variable in a logistic regression, where the number of IS inside and outside the tested gene (*y*=1) is subtracted from the length of the tested gene and the total length of the investigated genome (*y*=0). For analyses C) and D), we model clone size growth using logistic regression for proportion data as the response variable and include time in the set of covariates. In all analyses, we performed a likelihood ratio test for each gene. The test statistic is used to assign a p-value to each gene. The p-values are then corrected to account for multiplicity. In the simulation and experimental data analysis, the false discovery rate (FDR) correction method is used. All analyses can be done with an iterative approach, where significant genes are iteratively removed from the dataset until no significant genes are detected. For the clone fitness analyses (C) and (D), we implemented an optional robust statistical calculation to analyze the gene therapy clinical data. This approach was designed to reduce the likelihood of false positives— instances where significant gene-growth associations arise, not from mechanistic modifications of clone fitness due to the IS, but from clonal population dynamics. The clonal selection process, driven by the engraftment bottleneck and the need to maintain homeostasis, is expected to expand certain clones. These expansions are not caused by altered clone fitness due to IS location and are, therefore, likely to occur randomly across the genome. However, with a limited number of independent datasets (e.g., data from only one patient for β-thal and SCD), it becomes challenging to differentiate between mechanistic and clonal dynamics effects. To address this issue, we developed a procedure to reduce our false positive rate. For each gene, the robust statistic is calculated by excluding the IS with the highest count (summed across all cell types and time points) within that gene, mitigating the bias introduced by clonal population dynamics. The robust statistic serves as a filter, with the standard statistic reported only for genes that remain statistically significant when the robust statistic is applied.

### Simulation study

The simulation study focuses on chromosome 15, which is approximately 108 Mb in length. Five new genes, referred to as *TestGenes* (with length of 5, 10, 20, 40, 80 Kb), were added to the gene list at non-overlapping locations. The final list of 1032 genes, including the *TestGenes*, was used for simulating integrations across all simulations. The study explores various effect sizes (*h* = 1, 2, 4, 8, 16, 32, 64), where *h* = 1 represents the null hypothesis, and sample sizes *N* = 100, 200, 400, 800 IS for gene targeting analysis (A, B); and cumulative clone sizes *M* = 1000, 2000, 4000, 8000 for clone fitness analyses (C, D). For each combination of *h* and sample size, 1000 simulations were performed.

Gene targeting (A): IS were sampled without replacement, with probability proportional to 1 for IS outside the *TestGenes* and proportional to *h* for IS within *TestGenes*. Four datasets, each with *N* integrations, were generated for each simulation. Differential gene targeting (B): IS were sampled without replacement, with probability in *TestGenes* proportional to: *h* for IS within *TestGenes* in group 1, 1 for IS within *TestGenes* in group 2. For the 1,027 endogenous genes, the IS probability (same in both groups) was proportional to a value sampled from an exponential distribution with parameter 1. The baseline IS rate outside of genes was set at 0.001. Each simulation produced four datasets of size *N* for each group. Clone fitness (C): A total of 10,000 IS were generated using the method described for group 2 in analysis B, representing the total population of IS. At each time point, *M* elements from the total population are sampled (with replacement). IS clone sizes are defined by the times each IS is sampled. Selective growth for IS within *TestGenes* is simulated by increasing the sampling probability over time for IS within *TestGenes*, while for IS outside *TestGenes* remained constant. The un-normalized probability for an IS within *TestGenes* evolved as (*q_it_ = 1 + (h-1) t*), while it remained *q_it_ =* 1 for those outside. Differential clone fitness (D): Two sets of 10,000 IS were generated for group 1 and group 2, similar to (C). Clones were sampled for both groups at each time point, with IS outside *TestGenes* evolving with a random growth rate sampled from a uniform distribution [min= 0, max=2] for both groups. For *TestGenes*, group 1 had sampling probabilities evolving as *(q_it_ = 1 + h t)*, while group 2 followed *(q_it_ = 1 + t)*.

### Software implementation

MELISSA is available as R package at https://github.com/PellinLab/Melissa. Instructions for reproducing the results in this paper are available as a notebook in the same repository.

### Data availability

Raw data generated are available at with code PRJNA1090149.

## ACKNOWLEDGEMENT

This project has been funded in part with Federal funds from the National Heart, Lung, and Blood Institute, National Institutes of Health, Department of Health and Human Services, under Contract No. 75N92019D00018. Additional resources utilized include the Indiana University Melvin and Bren Simon Comprehensive Cancer Center Flow Cytometry Core (P30 CA082709) and the Sequencing analysis was carried out in the Center for Medical Genomics at Indiana University School of Medicine, which is partially supported by the Indiana University Grand Challenges Precision Health Initiative.

## AUTHOR CONTRIBUTIONS

T.Y.L., K.C. and D.P. conceived and designed the experiments and wrote the manuscript with input from all authors. D.P. designed the statistical method and G.C. software implementation and the simulation study. D.K. and C.B. performed HSPC transduction. T.Y.L., K.C, D.P. performed integration site analysis. T.Y.L., K.C., G.C., and D.P. provided analysis and interpretation.

