## supplementary tables for "Modeling integration site data for safety assessment with MELISSA"

| A) | p 01 | p 02 | p 04 | p 08 | p 16 | p 32 | p 64 | n 01 | n 02 | n 04 | n 08 | n 16 | n 32 | n 64 |
|---|---|---|---|---|---|---|---|---|---|---|---|---|---|---|
| 1h | 0.00 | 0.00 | 0.01 | 0.06 | 0.18 | 0.48 | 0.83 | 0.91 | 0.90 | 0.91 | 0.91 | 0.92 | 0.95 | 0.99 |
| | (0.03) | (0.06) | (0.12) | (0.24) | (0.38) | (0.49) | (0.34) | (0.03) | (0.03) | (0.03) | (0.04) | (0.04) | (0.05) | (0.03) |
| 2h | 0.00 | 0.00 | 0.01 | 0.05 | 0.27 | 0.72 | 0.96 | 0.97 | 0.97 | 0.98 | 0.98 | 0.99 | 1.00 | 1.00 |
| | (0.00) | (0.04) | (0.09) | (0.22) | (0.44) | (0.44) | (0.16) | (0.00) | (0.00) | (0.01) | (0.01) | (0.01) | (0.01) | (0.00) |
| 4h | 0.00 | 0.00 | 0.01 | 0.11 | 0.56 | 0.96 | 1.00 | 0.99 | 0.99 | 0.99 | 1.00 | 1.00 | 1.00 | 1.00 |
| | (0.00) | (0.04) | (0.11) | (0.31) | (0.49) | (0.18) | (0.04) | (0.00) | (0.00) | (0.00) | (0.00) | (0.00) | (0.00) | (0.00) |
| 8h | 0.00 | 0.00 | 0.01 | 0.24 | 0.86 | 0.99 | 0.99 | 1.00 | 1.00 | 1.00 | 1.00 | 1.00 | 1.00 | 1.00 |
| | (0.00) | (0.00) | (0.11) | (0.42) | (0.34) | (0.07) | (0.08) | (0.00) | (0.00) | (0.00) | (0.00) | (0.00) | (0.00) | (0.00) |

| B) | p 01 | p 02 | p 04 | p 08 | p 16 | p 32 | p 64 | n 01 | n 02 | n 04 | n 08 | n 16 | n 32 | n 64 |
|---|---|---|---|---|---|---|---|---|---|---|---|---|---|---|
| 1h | 0.00 | 0.00 | 0.00 | 0.00 | 0.00 | 0.01 | 0.41 | 0.99 | 0.99 | 0.99 | 1.00 | 1.00 | 1.00 | 1.00 |
| | (0.00) | (0.00) | (0.00) | (0.00) | (0.00) | (0.11) | (0.49) | (0.00) | (0.00) | (0.00) | (0.01) | (0.00) | (0.00) | (0.00) |
| 2h | 0.00 | 0.00 | 0.00 | 0.00 | 0.00 | 0.27 | 0.96 | 1.00 | 1.00 | 1.00 | 1.00 | 1.00 | 1.00 | 1.00 |
| | (0.00) | (0.00) | (0.00) | (0.00) | (0.03) | (0.44) | (0.17) | (0.00) | (0.00) | (0.00) | (0.00) | (0.00) | (0.00) | (0.00) |
| 4h | 0.00 | 0.00 | 0.00 | 0.00 | 0.18 | 0.94 | 0.98 | 1.00 | 1.00 | 1.00 | 1.00 | 1.00 | 1.00 | 1.00 |
| | (0.00) | (0.00) | (0.00) | (0.00) | (0.38) | (0.22) | (0.09) | (0.00) | (0.00) | (0.00) | (0.00) | (0.00) | (0.00) | (0.00) |
| 8h | 0.00 | 0.00 | 0.00 | 0.12 | 0.88 | 0.98 | 0.98 | 1.00 | 1.00 | 1.00 | 1.00 | 1.00 | 1.00 | 1.00 |
| | (0.00) | (0.00) | (0.00) | (0.32) | (0.31) | (0.09) | (0.18) | (0.00) | (0.00) | (0.00) | (0.00) | (0.00) | (0.00) | (0.00) |

| C) | p 01 | p 02 | p 04 | p 08 | p 16 | p 32 | p 64 | n 01 | n 02 | n 04 | n 08 | n 16 | n 32 | n 64 |
|---|---|---|---|---|---|---|---|---|---|---|---|---|---|---|
| 1k | 0.00 | 0.00 | 0.04 | 0.87 | 1.00 | 1.00 | 1.00 | 1.00 | 1.00 | 1.00 | 1.00 | 1.00 | 1.00 | 1.00 |
| | (0.00) | (0.00) | (0.19) | (0.34) | (0.03) | (0.00) | (0.00) | (0.00) | (0.00) | (0.00) | (0.00) | (0.00) | (0.00) | (0.00) |
| 2k | 0.00 | 0.00 | 0.79 | 1.00 | 1.00 | 1.00 | 1.00 | 1.00 | 1.00 | 1.00 | 1.00 | 1.00 | 1.00 | 1.00 |
| | (0.00) | (0.04) | (0.41) | (0.04) | (0.00) | (0.00) | (0.00) | (0.00) | (0.00) | (0.00) | (0.00) | (0.00) | (0.00) | (0.00) |
| 4k | 0.00 | 0.28 | 1.00 | 1.00 | 1.00 | 1.00 | 1.00 | 1.00 | 1.00 | 1.00 | 1.00 | 1.00 | 1.00 | 1.00 |
| | (0.00) | (0.45) | (0.07) | (0.00) | (0.00) | (0.00) | (0.00) | (0.00) | (0.00) | (0.00) | (0.00) | (0.00) | (0.00) | (0.00) |
| 8k | 0.00 | 0.91 | 1.00 | 1.00 | 1.00 | 1.00 | 1.00 | 1.00 | 1.00 | 1.00 | 1.00 | 1.00 | 1.00 | 1.00 |
| | (0.00) | (0.29) | (0.00) | (0.00) | (0.00) | (0.00) | (0.00) | (0.00) | (0.00) | (0.00) | (0.00) | (0.00) | (0.00) | (0.00) |

| D) | p 01 | p 02 | p 04 | p 08 | p 16 | p 32 | p 64 | n 01 | n 02 | n 04 | n 08 | n 16 | n 32 | n 64 |
|---|---|---|---|---|---|---|---|---|---|---|---|---|---|---|
| 1k | 0.00 | 0.00 | 0.00 | 0.00 | 0.00 | 0.00 | 0.17 | 1.00 | 1.00 | 1.00 | 1.00 | 1.00 | 1.00 | 1.00 |
| | (0.00) | (0.00) | (0.00) | (0.00) | (0.00) | (0.00) | (0.38) | (0.00) | (0.00) | (0.00) | (0.00) | (0.00) | (0.00) | (0.00) |
| 2k | 0.00 | 0.00 | 0.00 | 0.00 | 0.00 | 0.36 | 0.82 | 1.00 | 1.00 | 1.00 | 1.00 | 1.00 | 1.00 | 1.00 |
| | (0.00) | (0.00) | (0.00) | (0.00) | (0.00) | (0.48) | (0.38) | (0.00) | (0.00) | (0.00) | (0.00) | (0.00) | (0.00) | (0.00) |
| 4k | 0.00 | 0.00 | 0.00 | 0.02 | 0.66 | 0.93 | 0.98 | 1.00 | 1.00 | 1.00 | 1.00 | 1.00 | 1.00 | 1.00 |
| | (0.00) | (0.00) | (0.00) | (0.13) | (0.47) | (0.25) | (0.13) | (0.00) | (0.00) | (0.00) | (0.00) | (0.00) | (0.00) | (0.00) |
| 8k | 0.00 | 0.00 | 0.03 | 0.82 | 0.99 | 0.99 | 1.00 | 1.00 | 1.00 | 1.00 | 1.00 | 1.00 | 1.00 | 1.00 |
| | (0.00) | (0.00) | (0.17) | (0.39) | (0.10) | (0.09) | (0.05) | (0.00) | (0.00) | (0.00) | (0.00) | (0.00) | (0.00) | (0.00) |

**Suppl. Table 1: Proportion of positive predicted values and negative predicted values in simulated datasets:** A) for IS gene targeting; B) IS differential gene targeting; C) IS/clone fitness; D) IS/clone differential clone fitness. Columns 2 to 8 contain the mean and standard deviation (in brackets) for the positive predicted value of the various simulations. Columns 9 to 15 contain the mean and standard deviation for negative predicted values. The rows indicate the sample size.

| | s = 100 / s = 1000 | s = 200 / s = 2000 | s = 400 / s = 4000 | s = 800 / s = 8000 |
|---|---|---|---|---|
| A) gene target | 0.21 (0.09) | 0.22 (0.03) | 0.26 (0.04) | 0.35 (0.06) |
| B) diff. gene target | 0.24 (0.04) | 0.26 (0.03) | 0.31 (0.03) | 0.39 (0.06) |
| C) clone fitness | 0.68 (0.12) | 1.04 (0.14) | 1.75 (0.20) | 3.12 (0.28) |
| D) diff. clone fitness | 2.20 (0.19) | 4.18 (0.32) | 8.05 (0.34) | 14.84 (0.73) |

**Suppl. Table 2: Computation time:** for different analyses and sample size, constant effect size (h = 8). Values are average time measured in seconds across 30 simulations, with their associated standard deviations (also in seconds) in brackets. For analyses A) and B) the number of integrations of each dataset is (s = 100, 200, 400, 800). For A) 4 datasets are used, whereas for B) 4 datasets for each group are used (8 in total). For analyses C) and D) the sum of the clone sizes of each dataset is (s = 1000, 2000, 4000, 8000). For analysis C) 6 datasets are used (one for each time), whereas for analysis D) 6 datasets are used for each group (12 in total).

| Sample | Passage | Group | IScount | Cumulative Clone Size | Diversity (H-index) | Estimated Tot. Number of IS (chao1 Estimates) | Diversity (Simpson's D) | UC50 |
|---|---|---|---|---|---|---|---|---|
| **Adip_MSC_P1** | 1 | Adip_MSC | 5820 | 5995 | 8.66 | 129447.81 | 1 | 2823 |
| **Adip_MSC_P4** | 4 | Adip_MSC | 4067 | 5072 | 8.2 | 17221.76 | 1 | 1532 |
| **Adip_MSC_P6** | 6 | Adip_MSC | 2295 | 3867 | 7.39 | 6279.07 | 1 | 489 |
| **Adip_MSC_P8** | 8 | Adip_MSC | 1255 | 3149 | 6.48 | 3374.22 | 1 | 141 |
| **BM_MSC_P1** | 1 | BM_MSC | 3919 | 4052 | 8.26 | 71413.39 | 1 | 1894 |
| **BM_MSC_P4** | 4 | BM_MSC | 3998 | 4566 | 8.24 | 23589.06 | 1 | 1716 |
| **BM_MSC_P6** | 6 | BM_MSC | 4533 | 5759 | 8.32 | 15504.31 | 1 | 1654 |
| **BM_MSC_P8** | 8 | BM_MSC | 3086 | 6042 | 7.68 | 6552.85 | 1 | 591 |
| **CD34_HSPC_r1** | 1 | CD34_HSPC | 13955 | 15357 | 9.51 | 87841.36 | 1 | 6277 |
| **CD34_HSPC_r2** | 1 | CD34_HSPC | 7261 | 8554 | 8.82 | 31787.19 | 1 | 2985 |
| **CD34_HSPC_r3** | 1 | CD34_HSPC | 14529 | 16182 | 9.54 | 88832.72 | 1 | 6439 |

**Suppl. Table 3: IS datasets summary table.** Samples' specific descriptive statistics and diversity index estimates. IScount: total number of IS. Cumulative Clone Size: Cumulative clone size. Diversity (H-index and Simpson's D (Whittaker, 1972)): clone diversity estimates calculated using Shannon H- index and Simpson's D index. Estimated Tot. Number of IS (chao1 Estimates, seen+unseen IS): estimates for the total number of IS present in the sample (Chao1 method (Chao, 1987)). UC50: minimum number of unique IS contributing for at least 50% of the cumulative clone size (Berry et al., 2017).

| AccessionNumber | CellType | HistoneModification |
|---|---|---|
| GSM669920 | Bone_Marrow_Derived_MSC_Cultured_Cells | H3K4me1 |
| GSM669957 | Bone_Marrow_Derived_MSC_Cultured_Cells | H3K27me3 |
| GSM670019 | Bone_Marrow_Derived_MSC_Cultured_Cells | H3K4me3 |
| GSM670028 | Bone_Marrow_Derived_MSC_Cultured_Cells | H3K9me3 |
| GSM670037 | Bone_Marrow_Derived_MSC_Cultured_Cells | H3K36me3 |
| GSM621398 | Adipose_Derived_MSC_Cultured_Cells | H3K9me3 |
| GSM621420 | Adipose_Derived_MSC_Cultured_Cells | H3K27me3 |
| GSM772747 | Adipose_Derived_MSC_Cultured_Cells | H3K4me3 |
| GSM772748 | Adipose_Derived_MSC_Cultured_Cells | H3K4me1 |
| GSM772820 | Adipose_Derived_MSC_Cultured_Cells | H3K36me3 |
| GSM773041 | Mobilized_CD34_Primary_Cells | H3K4me3 |
| GSM773042 | Mobilized_CD34_Primary_Cells | H3K36me3 |
| GSM773043 | Mobilized_CD34_Primary_Cells | H3K4me1 |
| GSM773047 | Mobilized_CD34_Primary_Cells | H3K27me3 |

| | | |
|---|---|---|
| GSM773049 | Mobilized_CD34_Primary_Cells | H3K9me3 |

**Suppl. Table 4: Datasets used for the IS/epigenetic association analysis.**
