## supplementary figures for "Modeling integration site data for safety assessment with MELISSA"

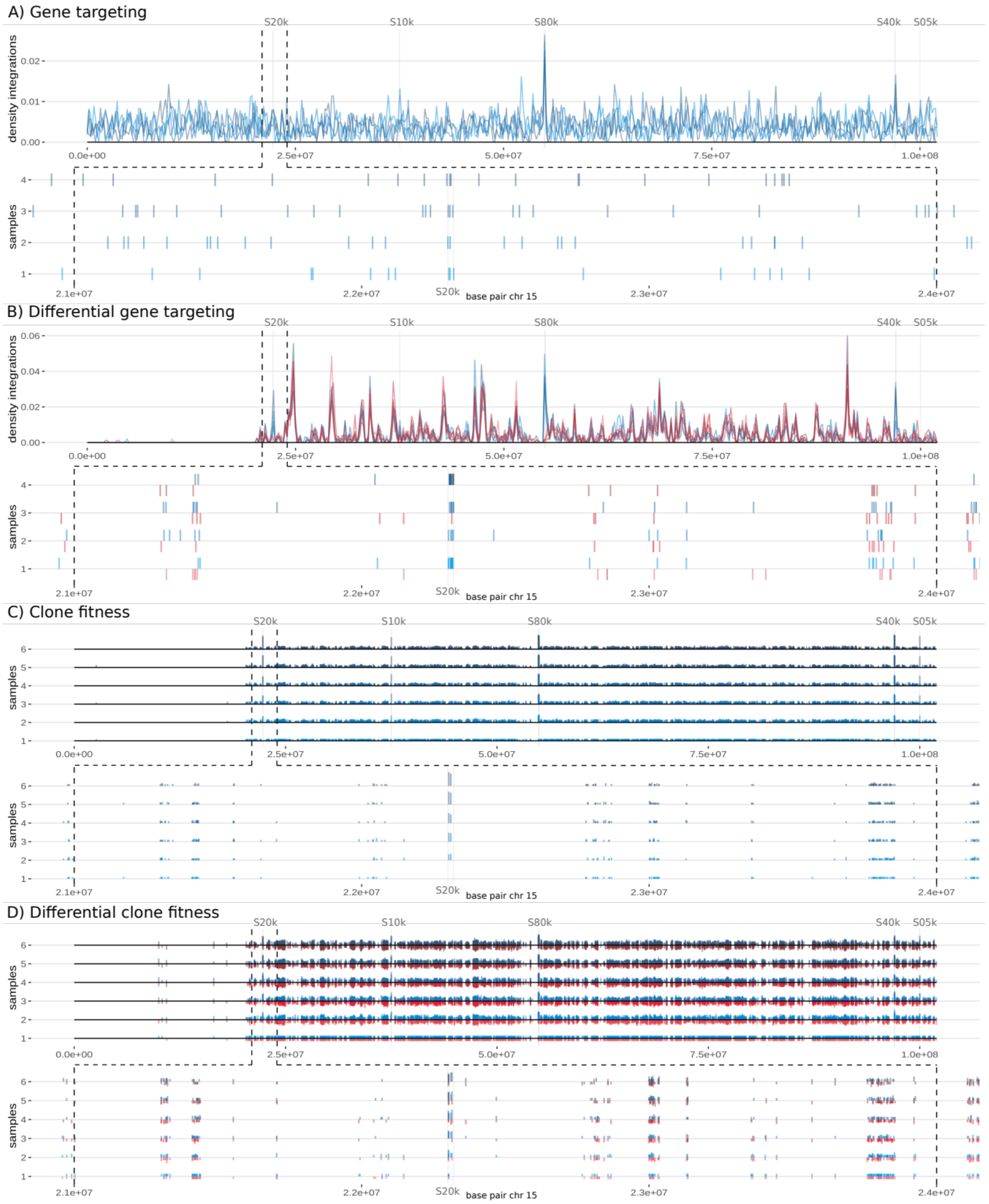


**Suppl. Figure 1: Example illustrating simulated datasets for models A), B), C), and D).** Each panel shows on the top the simulated IS datasets on chr15, whereas in the bottom part of each panel the genomic area surrounding a TestGene (length 20k, chr15: 2.23e07 - chr15: 2.232e07) is highlighted. The other TestGenes with lengths 5k, 10k, 80k, and 40k are also highlighted. For the differential analyses, the B) and D) values in blue are for group 1 and in red for group 2. The first plot for analyses A) and B) contains the kernel density of the IS for the four datasets. Below the IS in the four datasets are plotted in the highlighted segment. The number of IS for each dataset is ($N$ = 800), and the effect size is ($h$ = 16). For analyses C) and D), the clones are plotted above and below (in the highlighted segment) for all six datasets. The height of the bars corresponds to the clone sizes. The cumulative clone size for each dataset is ($M$ = 8000) and the effect size is ($h$ = 16).

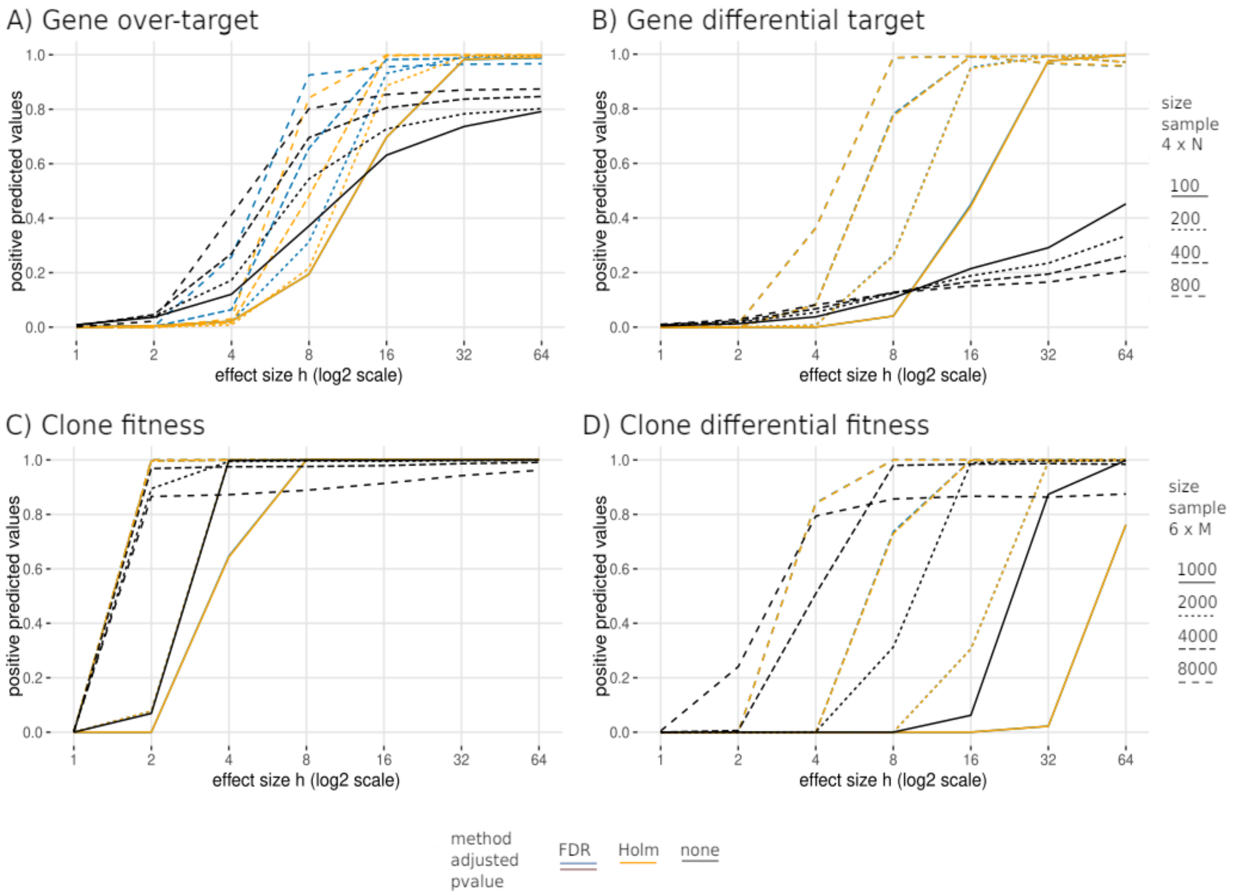


**Suppl. Figure 2: Impact of p-values adjustment criteria on the positive predicted values.** Average positive predicted values for model A) gene targeting rate; B) differential gene targeting; C) clone fitness; D) differential clone fitness applying False Discovery Rate, Holm method, and no correction (none). The results with FDR and Holm are almost identical, except for analysis A) where there are some minor differences between them.

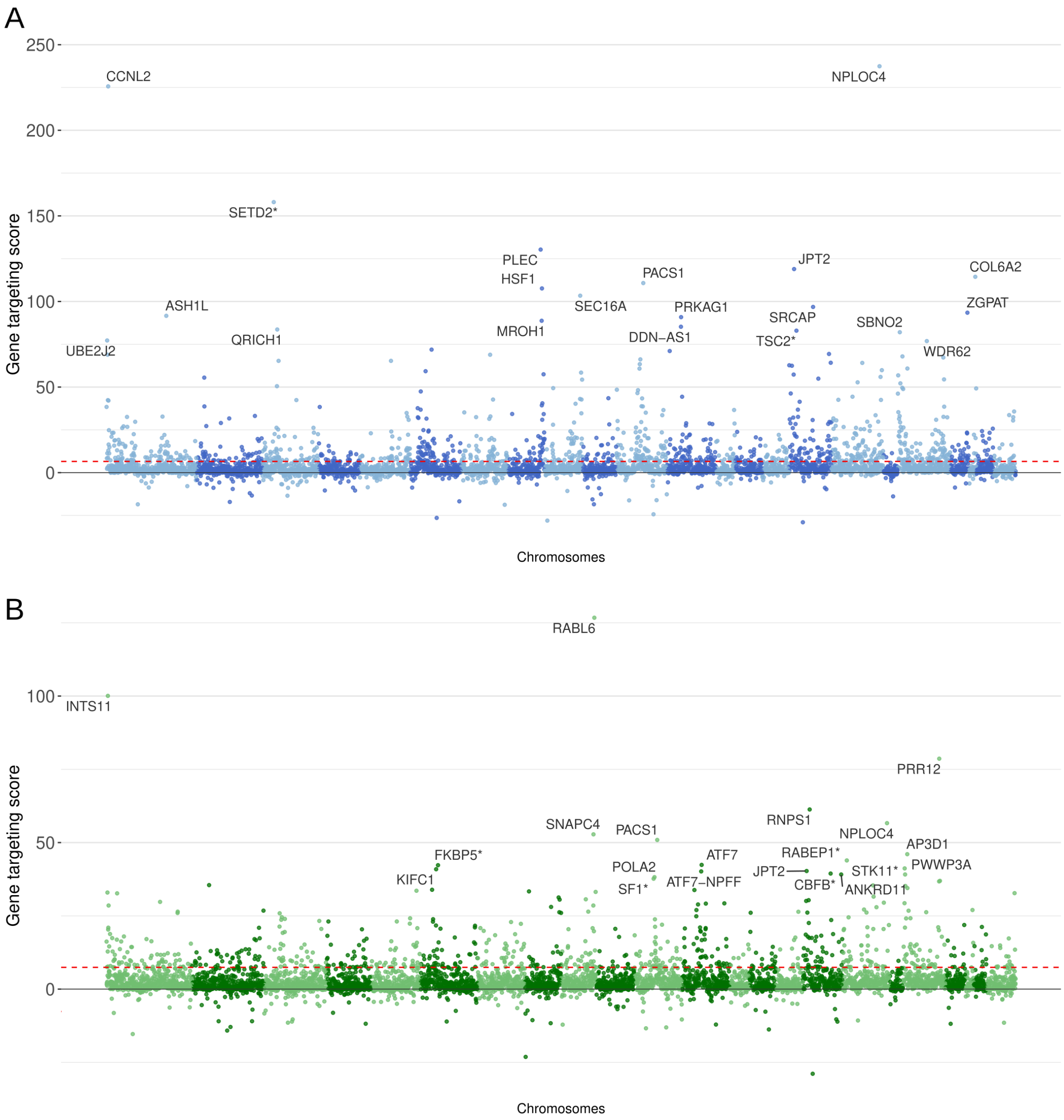


**Suppl. Figure 3: Gene targeting score for the A) BM MSC and B) Ad MSC dataset.** Each dot corresponds to a gene. On the x-axis, genes are ordered according to chromosome location, with genes in even-numbered chromosomes having a lighter color tone. The gene targeting score on the y-axis corresponds to the signed LRT test statistics. The dashed red line represents the threshold for statistically significant enrichment (LRT p-value adjusted with FDR, α=0.05).

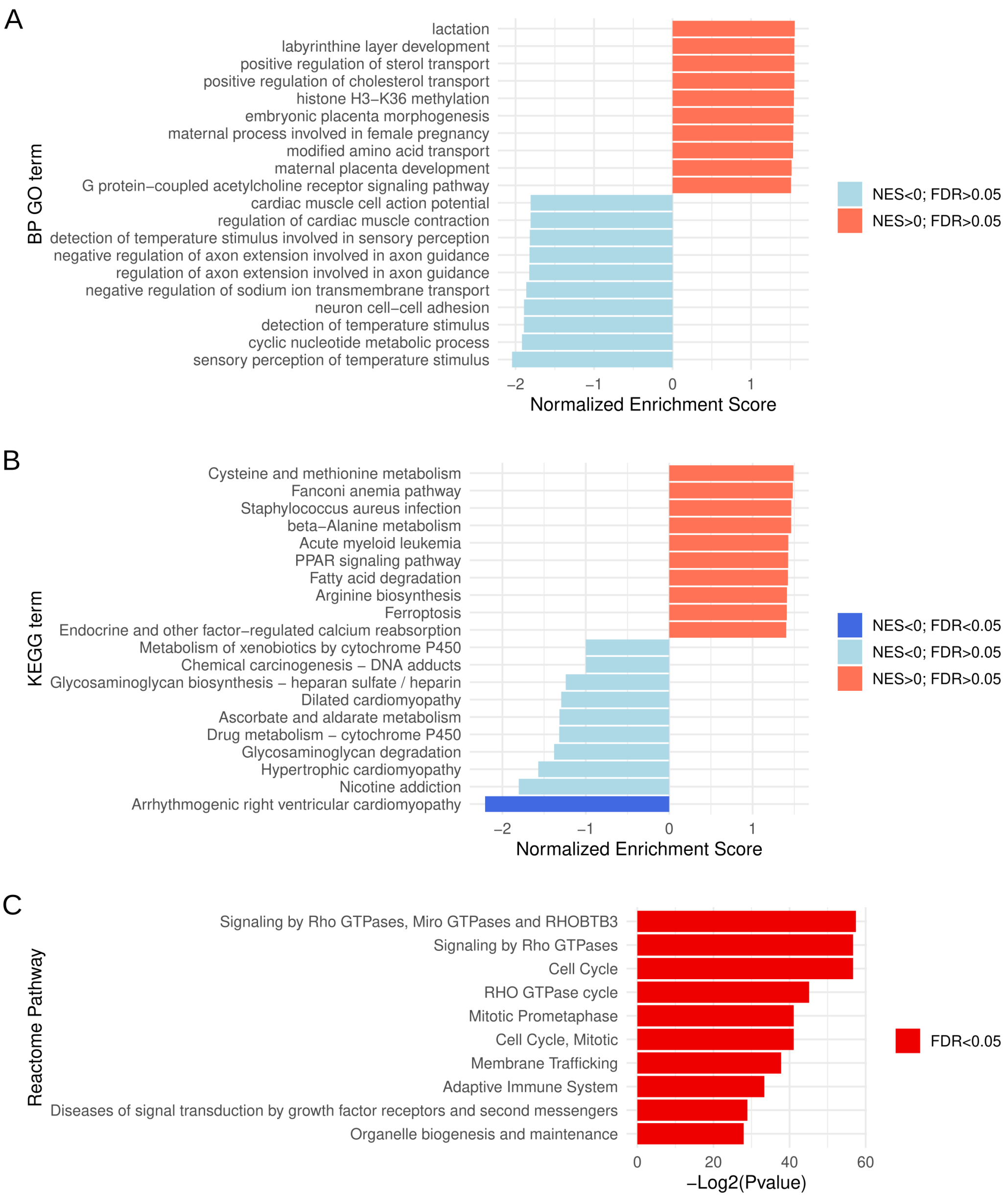


**Suppl. Figure 4: Pathway analysis on the gene targeting analysis on the HSPC dataset.** A) Gene Ontology enrichment analysis. B) KEGG pathway enrichment analysis. C) Reactome pathway analysis.

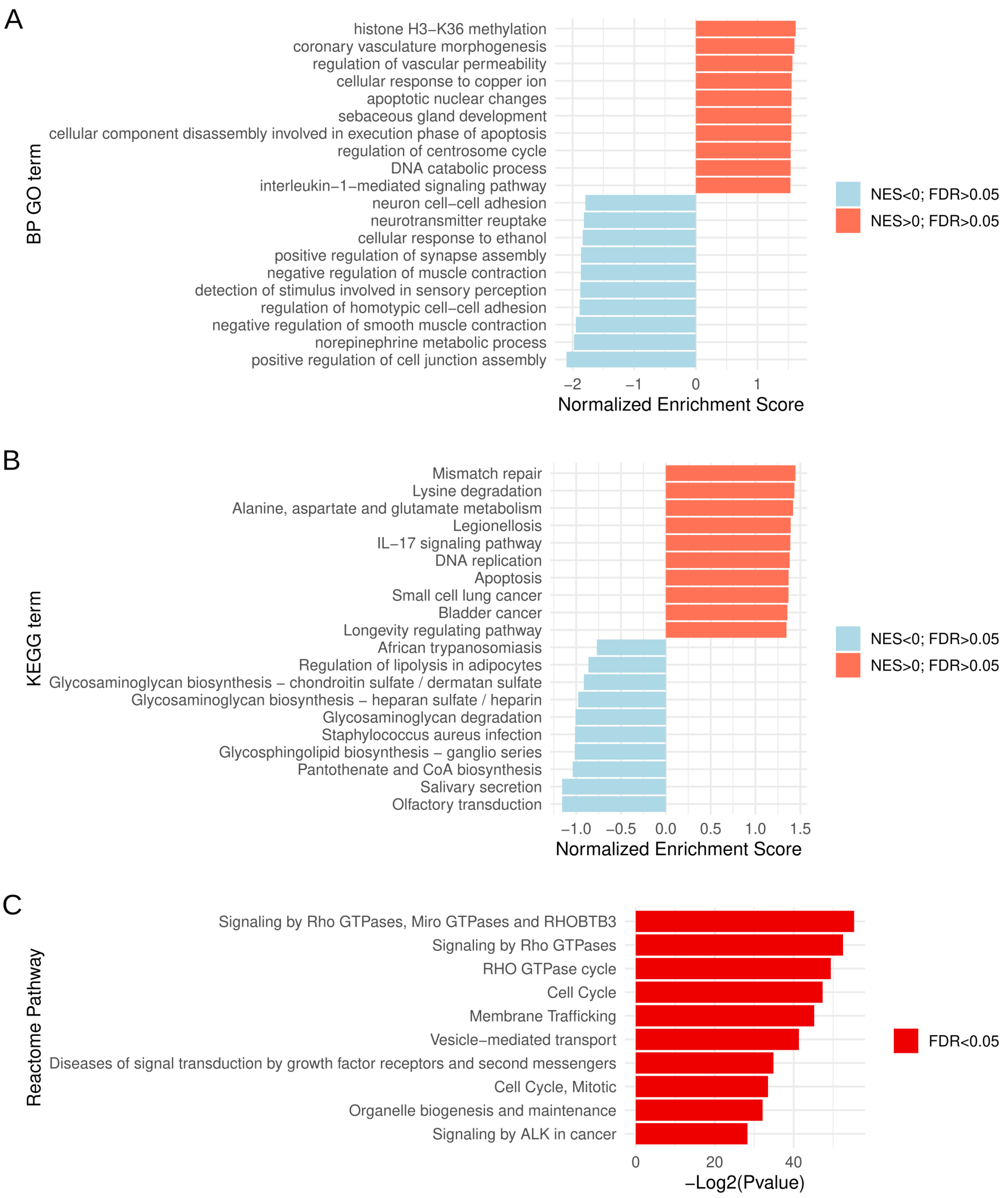


**Suppl. Figure 5: Pathway analysis on the gene targeting analysis on the BM MSC dataset.** A) Gene Ontology enrichment analysis. B) KEGG pathway enrichment analysis. C) Reactome pathway analysis.

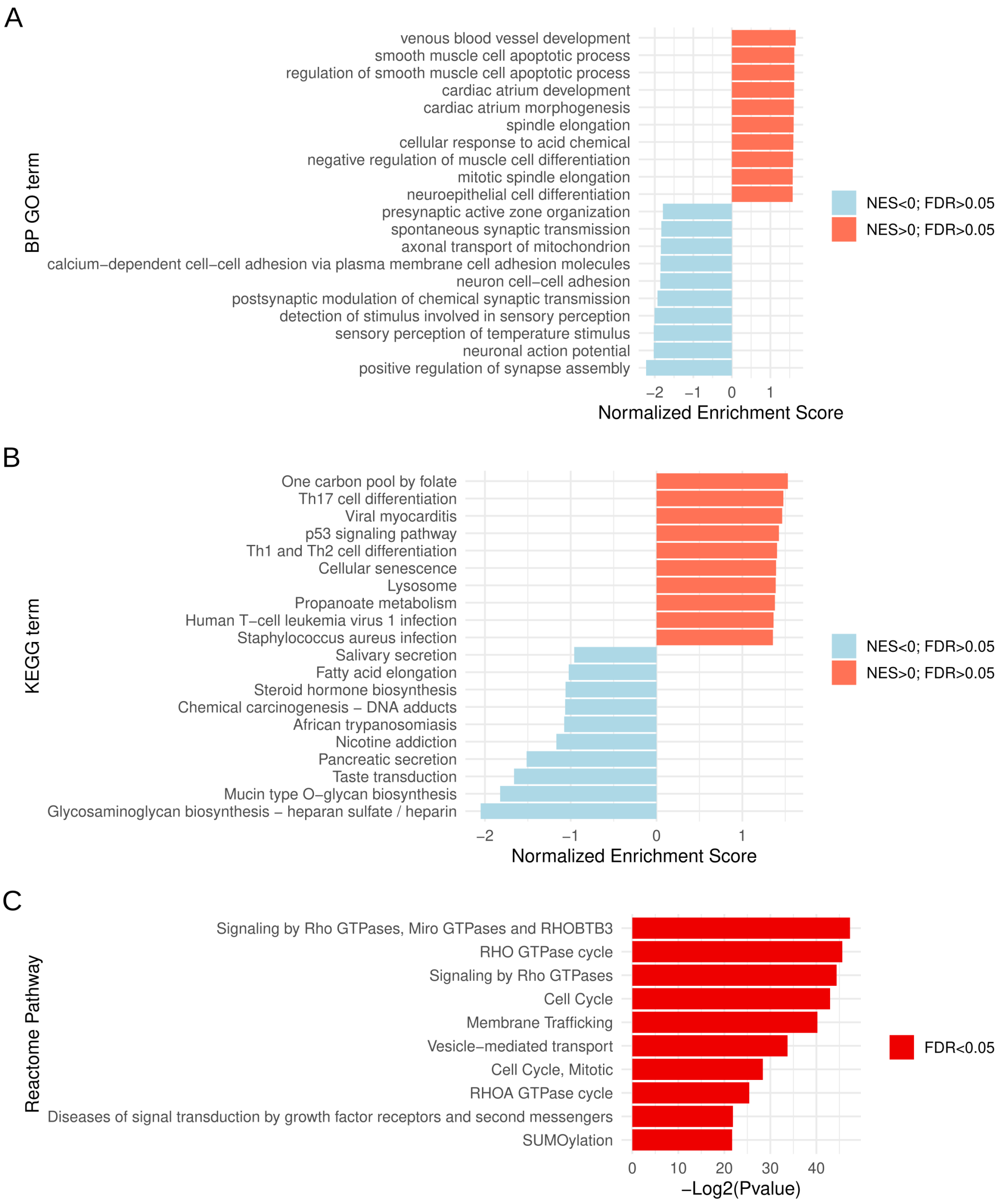


**Suppl. Figure 6: Pathway analysis on the gene targeting analysis on the Ad MSC dataset.** A) Gene Ontology enrichment analysis. B) KEGG pathway enrichment analysis. C) Reactome pathway analysis.

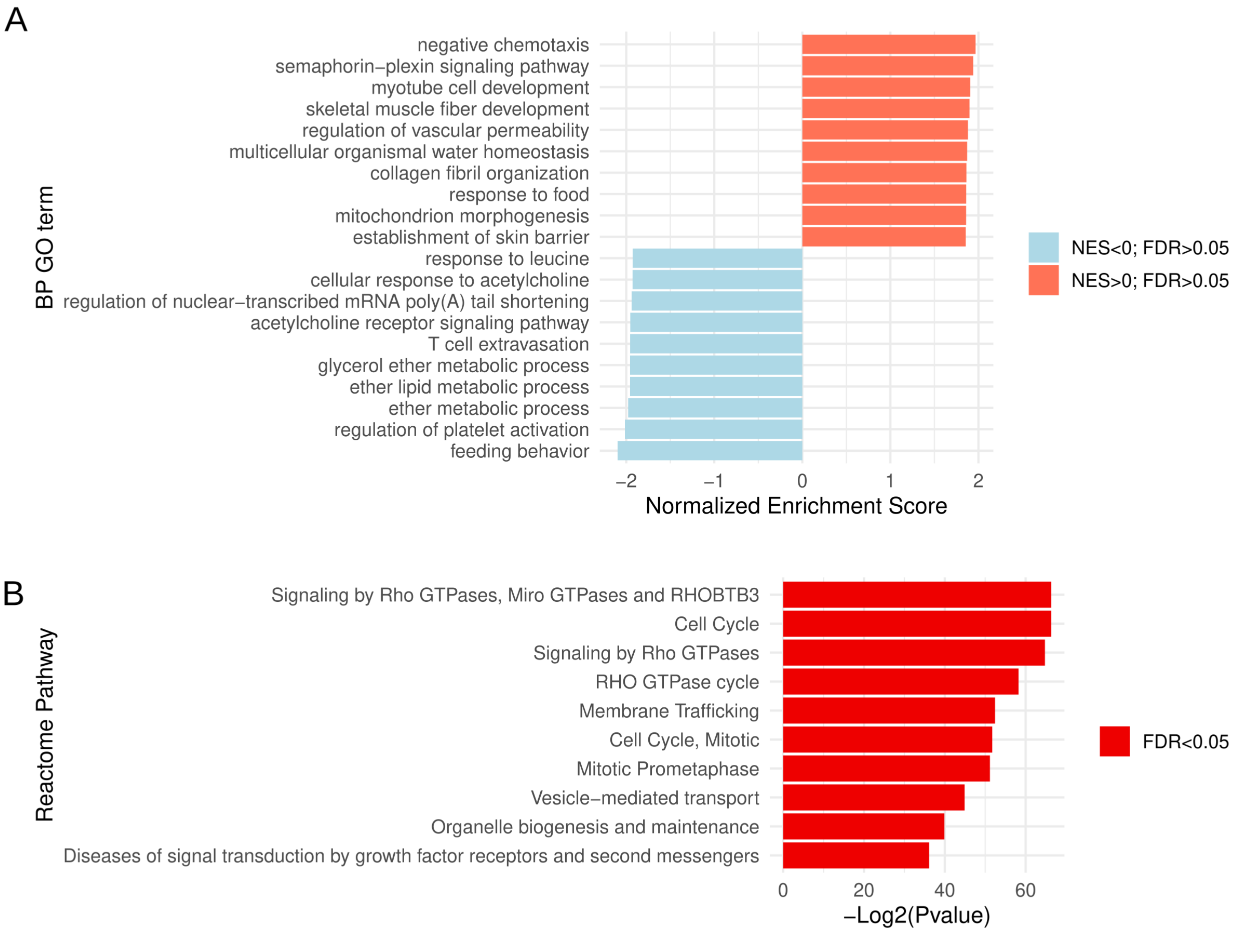


**Suppl. Figure 7: Pathway analysis on the differential gene targeting analysis comparing BM MSC and HSPC datasets.** A) Gene Ontology enrichment analysis. Positive (negative) NES corresponds to enrichment in BM MSC (HSPC). B) Reactome pathway analysis.

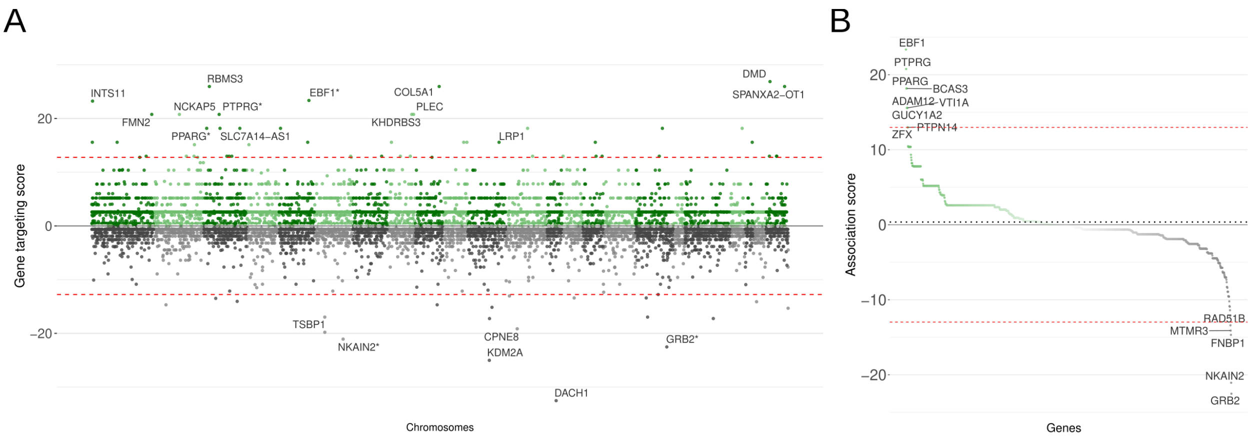


**Suppl. Figure 8: Ad MSC/HSPC differential gene targeting analysis.** A) Differential gene targeting analysis comparing Ad MSC and HSPC. Each dot represents a gene. If the gene is enriched in the integration site in Ad MSC (HSPC), it is plotted in the green (grey) section. Genes marked with a * are included in the high-risk gene list. The dashed red line represents the threshold for statistically significant enrichment. B) Ad MSC/HSPC waterfall plot for high-risk genes. Integration site enrichment scores for genes included in the high-risk genes are ranked by targeting score. The dashed red line represents the threshold for statistically significant enrichment (LRT p-value adjusted with FDR, α=0.05).

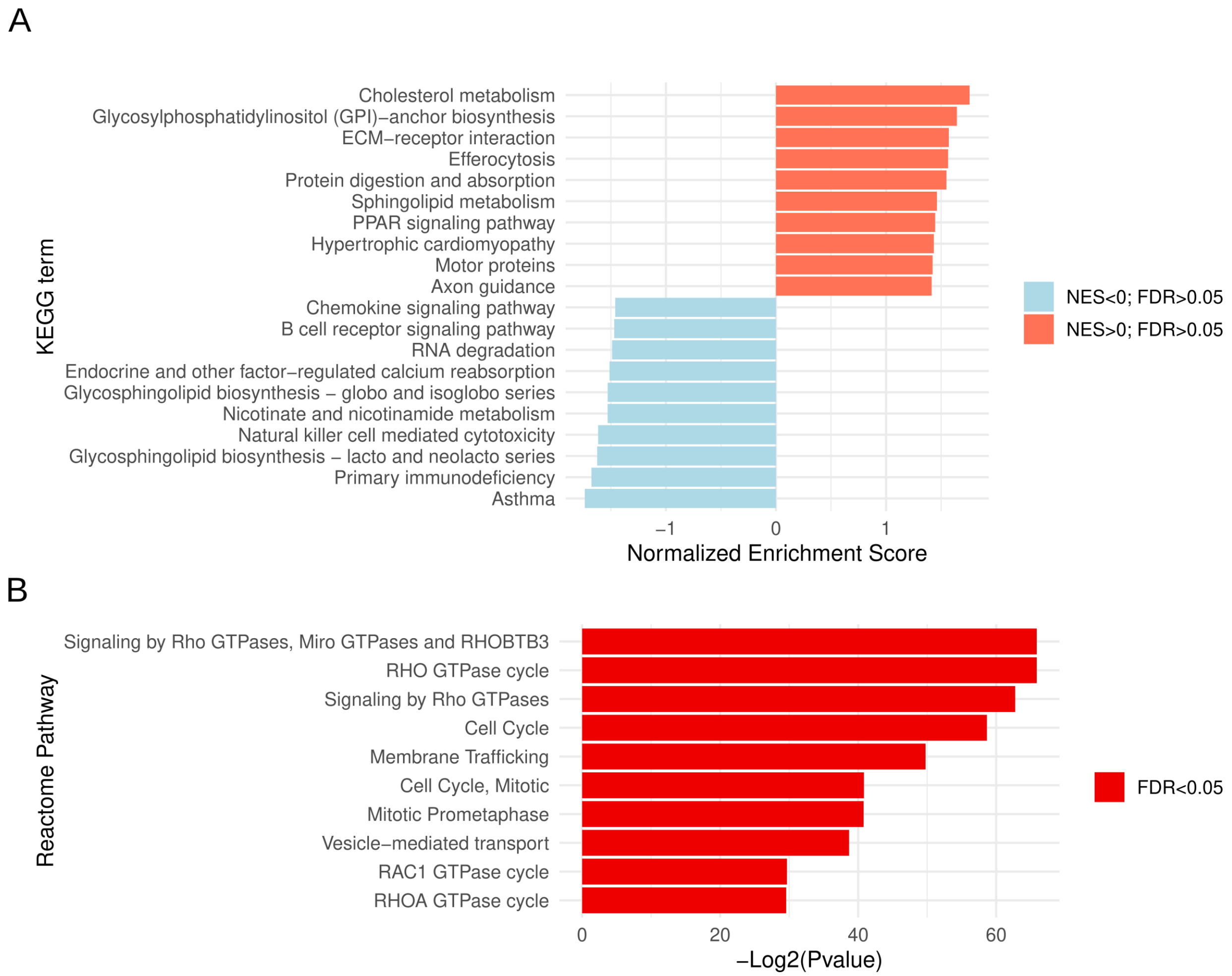


**Suppl. Figure 9: Pathway analysis on the differential gene targeting analysis comparing Ad MSC and HSPC datasets.** A) KEGG pathway enrichment analysis. Positive (negative) NES corresponds to enrichment in Ad MSC (HSPC). B) Reactome pathway analysis.

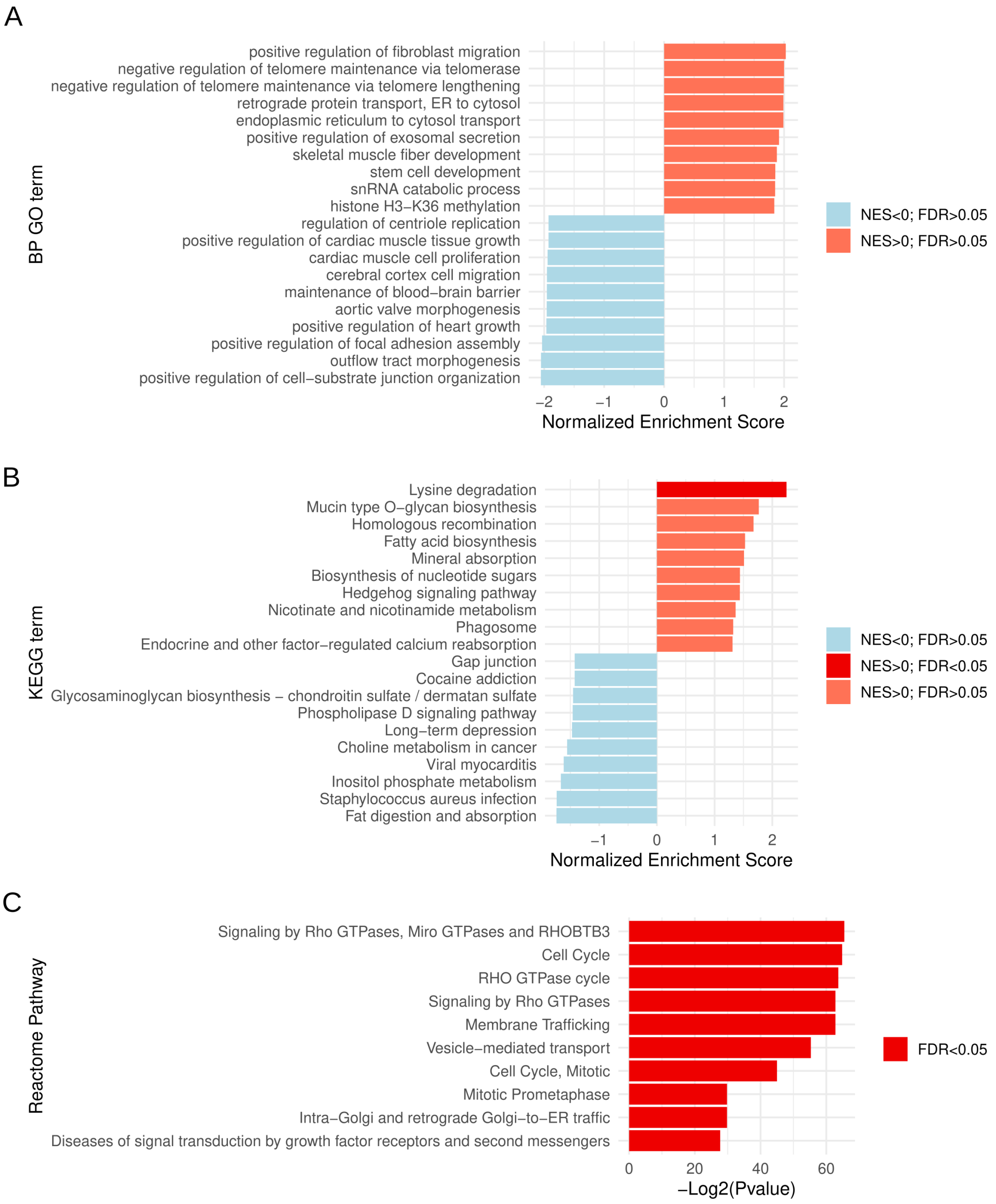


**Suppl. Figure 10: Pathway analysis on the differential gene targeting analysis comparing BM MSC and Ad MSC datasets.** A) Gene Ontology enrichment analysis. Positive (negative) NES corresponds to enrichment in BM MSC (Ad MSC). B) KEGG pathway enrichment analysis. Positive (negative) NES corresponds to enrichment in BM MSC (Ad MSC). C) Reactome pathway analysis.

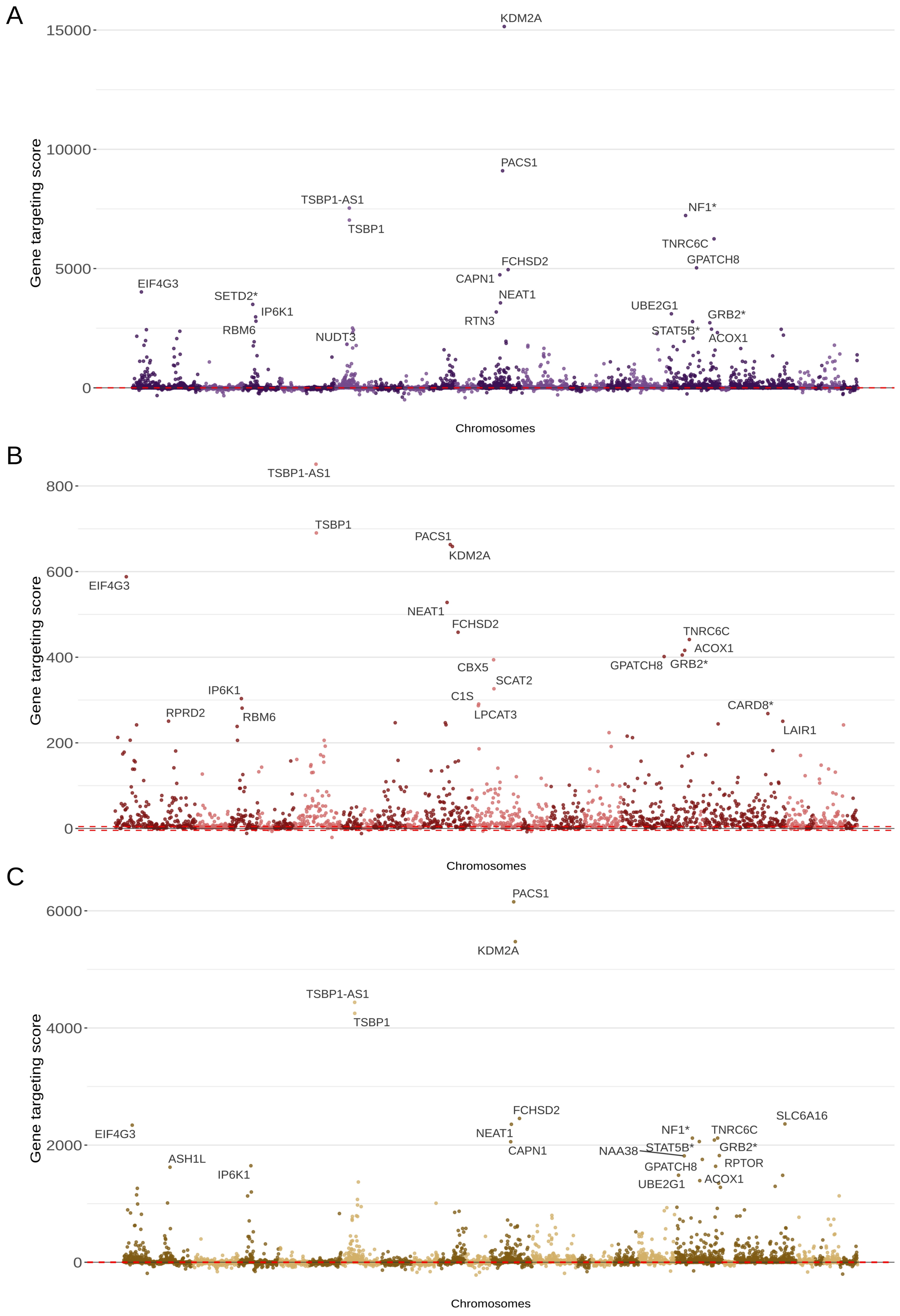

A
15000
10000
5000
0
Gene targeting score
KDM2A
PACS1
TSBP1-AS1
TSBP1
NF1*
TNRC6C
GPATCH8
FCHSD2
CAPN1
EIF4G3
SETD2*
IP6K1
NEAT1
UBE2G1
GRB2*
RTN3
RBM6
STAT5B*
NUDT3
ACOX1
Chromosomes
B
800
600
400
200
0
Gene targeting score
TSBP1-AS1
TSBP1
PACS1
KDM2A
EIF4G3
NEAT1
FCHSD2
TNRC6C
ACOX1
GPATCH8
GRB2*
CBX5
SCAT2
IP6K1
C1S
RPRD2
RBM6
LPCAT3
CARD8*
LAIR1
Chromosomes
C
6000
4000
2000
0
Gene targeting score
PACS1
KDM2A
TSBP1-AS1
TSBP1
FCHSD2
SLC6A16
EIF4G3
NEAT1
NF1*
TNRC6C
CAPN1
NAA38
STAT5B*
GRB2*
ASH1L
GPATCH8
RPTOR
IP6K1
UBE2G1
ACOX1
Chromosomes

**Suppl. Figure 11:** Gene targeting score for the A) β-thal, B) SCD, and C) WAS clinical trial. Each dot corresponds to a gene. On the x-axis, genes are ordered according to chromosome location, with genes in even-numbered chromosomes having a lighter color tone. The gene targeting score on the y-axis corresponds to signed LRT test statistics. Genes marked with a * are included in the high-risk gene list. The dashed red line represents the threshold for statistically significant enrichment (LRT p-value adjusted with FDR, $\alpha$=0.05).

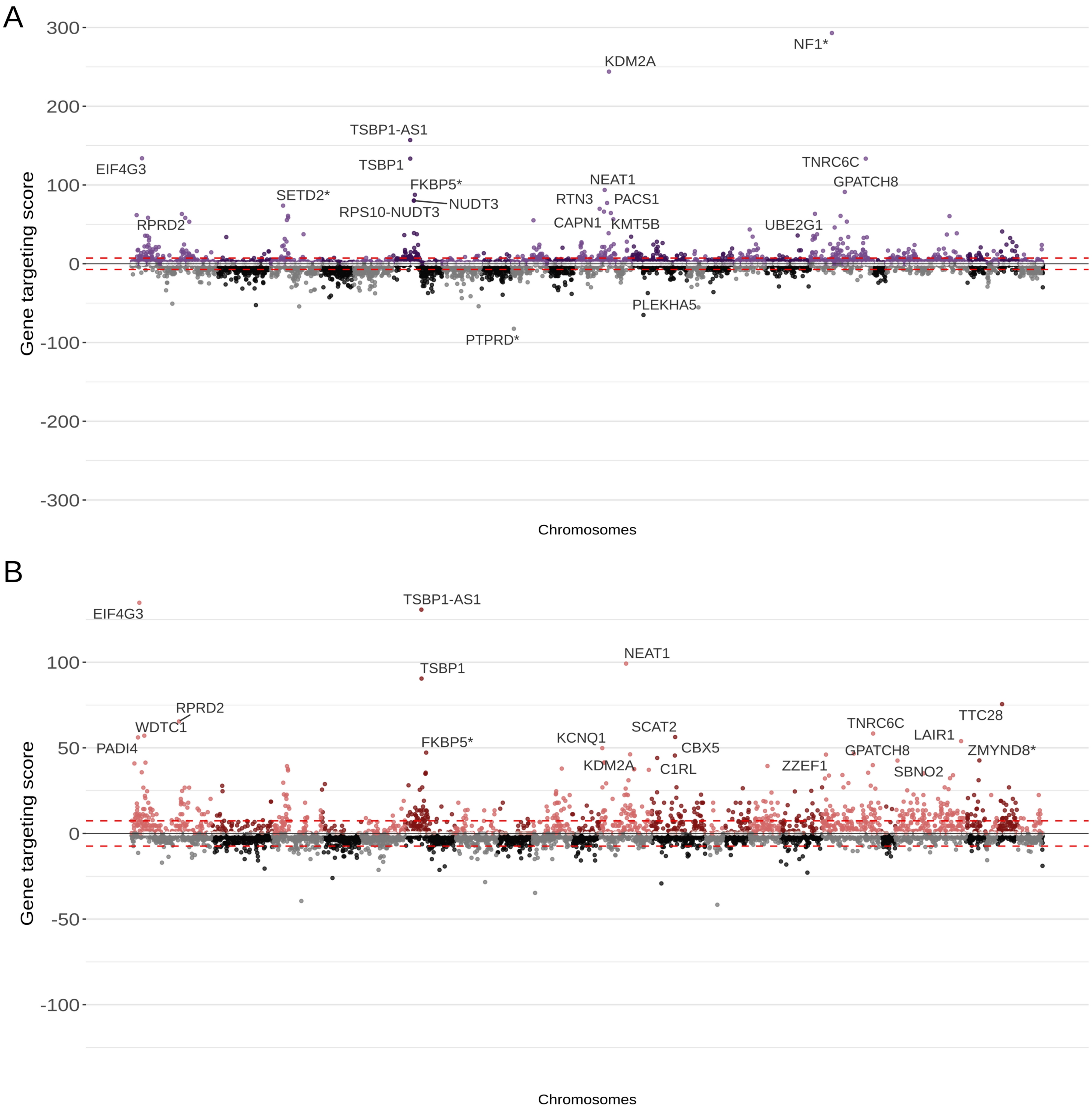


**Suppl. Figure 12: Differential gene targeting analysis on the clinical trails data.** Differential gene targeting analysis comparing A) β-thal and HSPC, B) SCD and HSPC. Each dot represents a gene. If the gene is enriched in the integration site in clinical trial datasets β-thal and SCD (HSPC), it is plotted in top (grey) section. Genes marked with a * are included in the high-risk gene list. The dashed red line represents the threshold for statistically significant enrichment (LRT p-value adjusted with FDR, α=0.05).

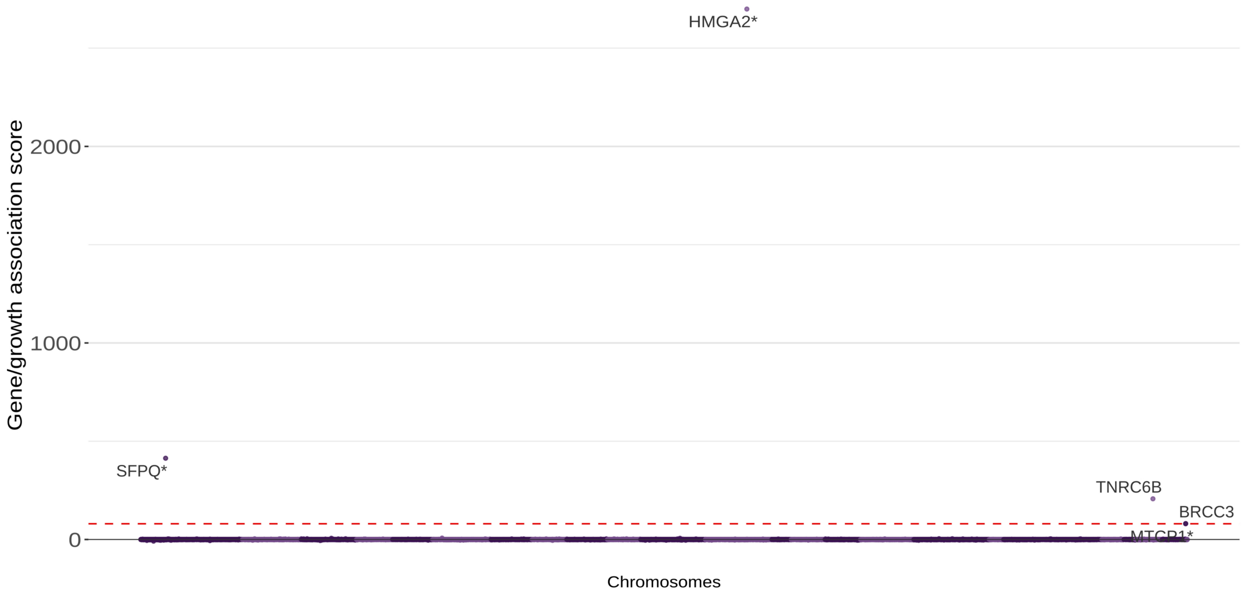


**Suppl. Figure 13:** Gene/growth association in the β-thal trial. The gene/growth association score provides a metric for assessing the significance of the fitness advantage or disadvantage conferred by IS within a gene. Genes marked with a * are included in the high-risk gene list. The dashed red line represents the threshold for statistically significant differential clone dynamics (LRT p-value adjusted with FDR, $\alpha=0.05$).

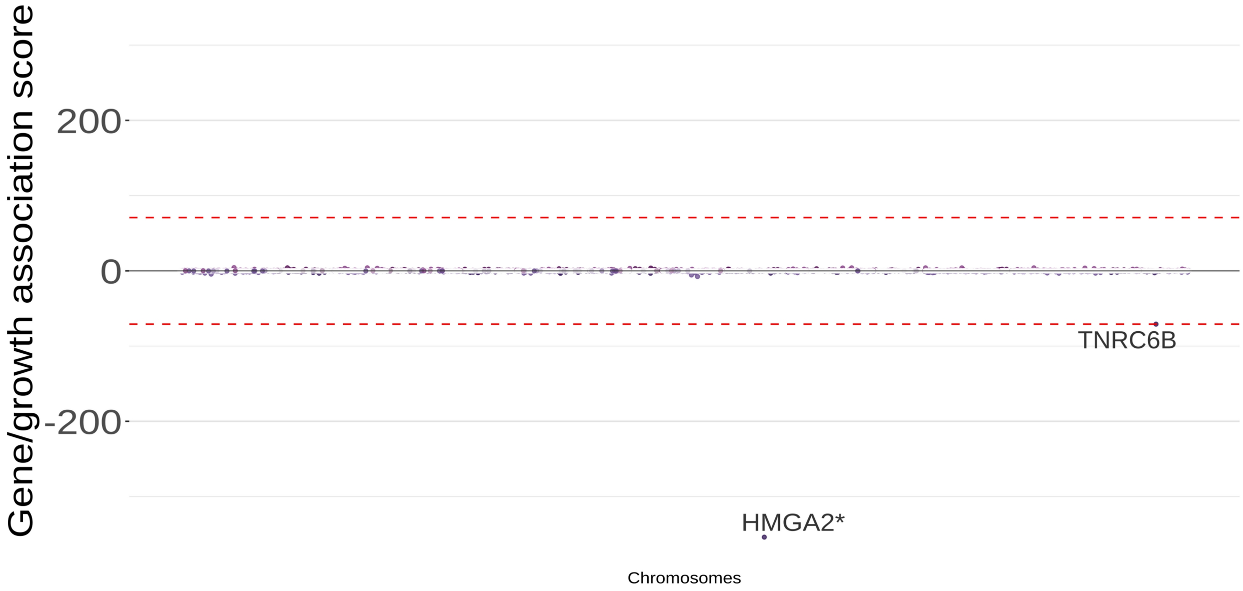


**Suppl. Figure 14: Lineage specific differential clone fitness analysis on β-thal trial data.** Enrichment score for differential clone growth dynamics in Lymphoid and Myeloid lineages in the β-thal. The plot illustrates the growth association score, quantifying the differential impact of integration site targeting genes on the growth dynamics of clones. If IS in the gene increases clone growth in Lymphoid (Myeloid), it is plotted in the top (bottom) section. Genes marked with a * are included in the high-risk gene list. The dashed red line represents the threshold for statistically significant differential clone dynamics (LRT p-value adjusted with FDR, α=0.05).
